# Modeling signaling-dependent pluripotent cell states with boolean logic can predict cell fate transitions

**DOI:** 10.1101/115683

**Authors:** Ayako Yachie-Kinoshita, Kento Onishi, Joel Ostblom, Eszter Posfai, Janet Rossant, Peter W. Zandstra

## Abstract

Pluripotent stem cells (PSCs) exist in multiple stable states, each with specific cellular properties and molecular signatures. The process by which pluripotency is either maintained or destabilized to initiate specific developmental programs is poorly understood. We have developed a model to predict stabilized PSC gene regulatory network (GRN) states in response to combinations of input signals. While previous attempts to model PSC fate have been limited to static cell compositions, our approach enables simulations of dynamic heterogeneity by combining an Asynchronous Boolean Simulation (ABS) strategy with simulated single cell fate transitions using Strongly Connected Components (SCCs). This computational framework was applied to a reverse-engineered and curated core GRN for mouse embryonic stem cells (mESCs) to simulate responses to LIF, Wnt/β-catenin, FGF/ERK, BMP4, and Activin A/Nodal pathway activation. For these input signals, our simulations exhibit strong predictive power for gene expression patterns, cell population composition, and nodes controlling cell fate transitions. The model predictions extend into early PSC differentiation, demonstrating, for example, that a Cdx2-high/Oct4-low state can be efficiently and robustly generated from mESCs residing in a naïve and signal-receptive state sustained by combinations of signaling activators and inhibitors.

**One Sentence Summary:** Predictive control of pluripotent stem cell fate transitions

## INTRODUCTION

Single cell-level heterogeneity in gene expression is common in pluripotent stem cells (PSCs)^1^ (and indeed other stem cell types^2^). There are two scenarios from which this heterogeneity emerges. Either different closely related cell types coexist, or individual cells transition dynamically between different cell states^3–5^. This diversity results in families of gene regulatory networks (GRNs), each with potentially unique responsiveness to endogenous or exogenous perturbations^6–8^. One manifestation of this is that different subpopulations of cells have higher probabilities of generating specific types of differentiated cells following treatment with differentiation-inducing ligands^9,10^.

Distinct PSCs and their associated GRNs appear to be stabilized through extrinsic signals (or signal modifiers)^11^, which are typically either supplemented into the medium or endogenously produced^12,13^. For example, mESCs cultured in LIF and BMP4, upon replacement with bFGF and Activin A, transition into epiblast stem cells (EpiSCs)^14,15^. Additionally, dual small molecule inhibition of MEK and glycogen synthase kinase-3ß (GSK3ß) (referred to as the 2i condition) drives mESCs back into naïve/ground state pluripotency, a state which closely resembles early, pre-implantation stage epiblast^16–18^. GRNs themselves do not only serve as responsive elements to external stimuli, but also as stimulus sources themselves (via autocrine/paracrine signaling), resulting in combined endogenous/exogenous feedback loops^13^ that influence cell fate transition probabilities.

Here we describe a simulation framework that quantitatively depicts each PSC subpopulation as a compilation of heterogeneous gene expression profiles. The computational framework uses a pruned mESC GRN consisting of 29 key genes to simulate regulation of Oct4, Sox2, and Nanog (as well as other mESC-associated genes) as a function of signaling inputs. Existing Boolean models that simulate the regulation of PSC GRNs^19–21^ treat each subpopulation as discrete, steady-state gene expression profiles (i.e. attractors) derived from unique or randomly set initial profiles after rounds of Boolean updates, where genes are toggled on and off to satisfy the Boolean logic that scaffolds the GRN. While the steady-state attractor approach simulates the presence of different cell states (i.e. subpopulations) within a total PSC population, it does not simulate gene expression variability within each PSC subpopulation. Importantly, these approaches do not capture observations gathered from single cell transcriptome data wherein variability^1,21^ and dynamics^22,23^ of gene expression have been observed between individual cells within a subpopulation. We therefore aimed to group closely related gene expression profiles into constructs that enabled the prediction of subpopulation composition. To do this we used an asynchronous Boolean simulation (ABS) strategy. While in a synchronous Boolean paradigm all genes in the GRN toggle simultaneously to produce each condition, ABS toggles individual genes asynchronously, resulting in a wider catalogue of transitional expression profiles^24^. Instead of depicting subpopulations as steady-state attractors, we account for transitional cell states by organizing transitional gene expression profiles into groups with strongly connected components (SCC) forming a closed loop between profiles (Figure S1a). Uniquely, this methodology allows the simulation outputs to be quantitatively compared with experimental observations from both single cell- and population-level experiments (Figure S1b.). This strategy more closely depicts the underlying biology of dynamic GRNs. In addition, our model also includes feedback loops from the GRN to signaling pathway components, thus allowing for the exploration of a broader and more nuanced array of GRN outputs such as the exit from pluripotency as a consequence of the activation and inhibition of different combinations of five major pluripotency-related signaling pathways (LIF/pStat3, Wnt/β-catenin, Bmp4/pSmad1/5/8, Activin A/pSmad2/3 and bFGF/pERK).

## RESULTS

### Simulation framework for PSCs

Our simulation framework took advantage of two key strategies. The first is the ABS strategy, where Boolean logical updates are performed asynchronously on individual nodes (genes) in each simulation update, allowing for multiple, unique transitional gene expression profiles to be generated from each single input profile^24,25^. A profile transition graph, the accumulation of the transitions between unique expression profiles, is analogous to transitions between single cell states and is derived from an iterative, random ABS (R-ABS). The relative transition frequencies from one expression profile to its successor profiles can be calculated by counting the individual transitions from the source to the target, which in turn determines the probability of the existence of each profile.

After generation of the profile transition graph through R-ABS, the second key element of our approach is to use SCC to group unique expression profiles. We define an SCC as the subset of expression profiles where every profile is capable of transitioning into all other profiles in the subset and returning to the original profile over an indefinite number of Boolean updates. This is analogous to a sustained PSC population containing multiple dynamically transitioning subpopulations^31^. In the context of population-level PSC state transitions, SCCs represent a dynamically stabilized population as a cluster of heterogeneous single cell profiles where each cell state can potentially give rise to any of the other states within the population. The model outputs provide predictions of the emergence of subpopulations (SCCs) in response to different input conditions. The gene expression level for any given gene within a particular SCC is predicted by multiplying the summation of expression profile probabilities where the gene is present (ON) by the sum of probabilities of all the profiles in the SCC and by further subtracting the probabilities of the out-going transitions (i.e. GRN profiles produced in the SCC that cannot give rise to all other cell states in the population) (Fig 1, see Online Methods and Supplementary Notes section 1). The population-averaged gene expression level is thereby calculated by summation of these values within each SCC, which is weighted by the proportion of sub-populations (the number of unique profiles in each SCC).

**Figure 1.**
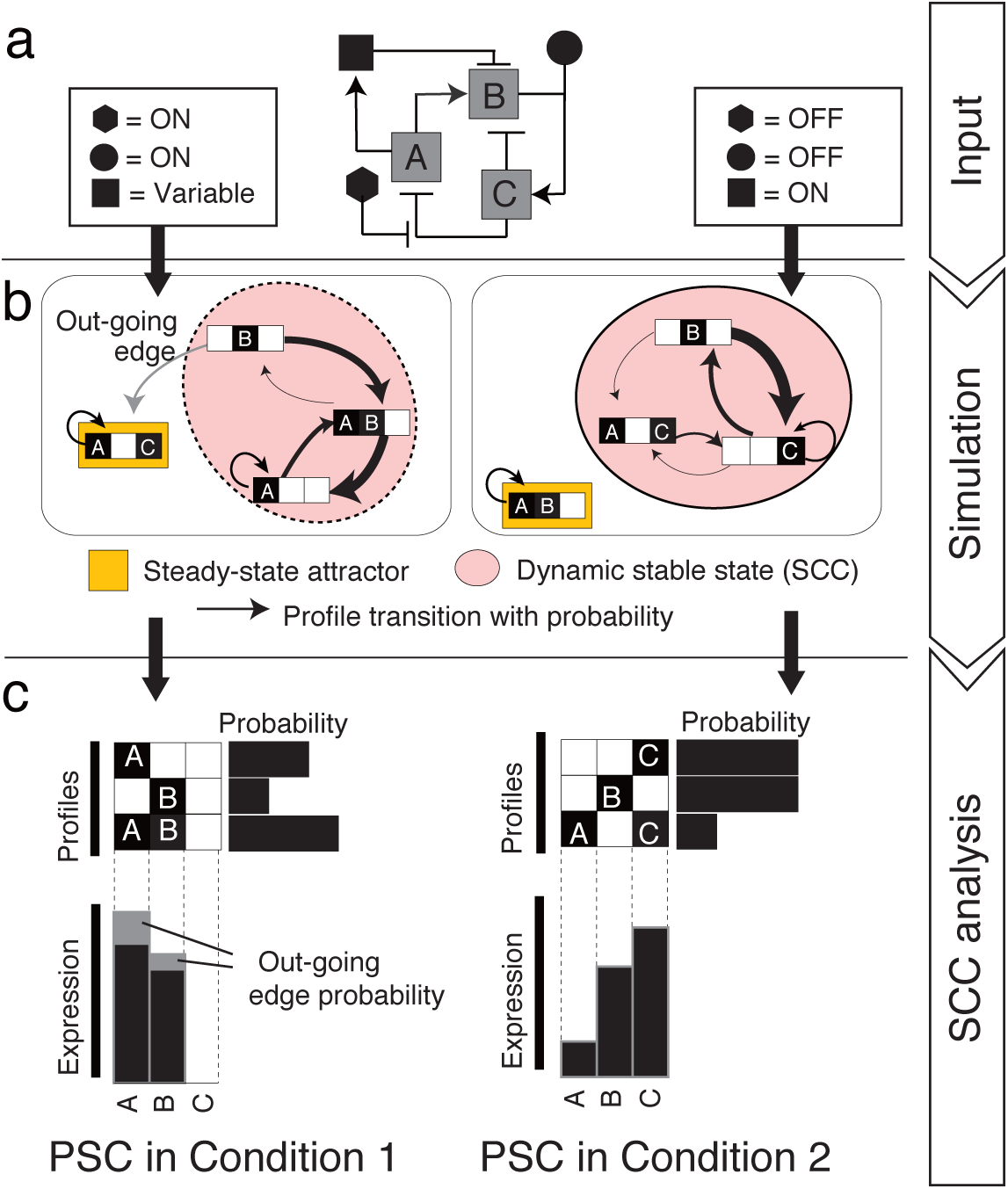
Strategy for modeling and simulation of stabilized PSCs. **(a)** Experimental conditions can be set as simulation inputs by defining the model variables (signal components with black-filled symbols and genes A, B and C) as either continuously ON, OFF, or variable. **(b)** Random ABS generates a directed transition graph where binary gene expression profiles are graph nodes and possible transitions from individual profiles are edges with a certain probability. PSC populations are assumed to be stabilized as a group of heterogeneous profiles, which is defined as an SCC. **(c)** Weighted, subpopulation-average gene expression and signaling activity of a particular SCC are calculated based on transition probabilities and the binary state of each model component in each heterogeneous profile.

### Mouse ESC-GRN construction

To build our GRN (see Supplementary Notes section 2 and 3 for details on GRN reconstruction), we first selected 14 pluripotency-specific genes based on prior knowledge^26,27^(Oct4, Sox2, Nanog, Klf4, c-Myc, Esrrb, Tbx3, Klf2, Gbx2, Jarid2, Mycn, Lrh1, Pecam1 and Rex1(Zfp42)), and key lineage specifiers known to drive the exit from pluripotency^28,29^(Tcf3, Cdx2, Gata6, Gcnf). EpiSC-enriched transcription factors (TFs), Fgf5, Eomes, Otx2, and Brachyury (T), were aggregated into a single component in the model termed EpiSC-enriched transcription factors (EpiTFs) for computational efficiency^14,30^. We then specified regulatory relationships among the genes by curation, including 19 regulations encompassing double positive or double negative regulatory circuits and known self-activations for seven genes (Supplementary Notes Table M1).

We next set the key signaling pathways (LIF/pStat3, Wnt/β-catenin, Bmp4/pSmad1/5/8, Activin A/pSmad2/3, bFGF/pERK and PI3K) as consequential effects of gene ON/OFF states by extending the gene list to link to signaling activities (Fgf4, Fgfr2, Bmp4 and Activin A/Nodal). For example, to model FGF activity and downstream MEK/ERK activation, we included stimulus sources (bFGF or Fgf4) and receptor availability (Fgfr2) in our GRN. To complete signal pathway integration, we then defined regulatory edges from cytokine/receptor-level signaling to their downstream effectors and feedback from genes to relevant signaling activities (Fig 2a and Supplementary Table S1.).

**Figure 2.**
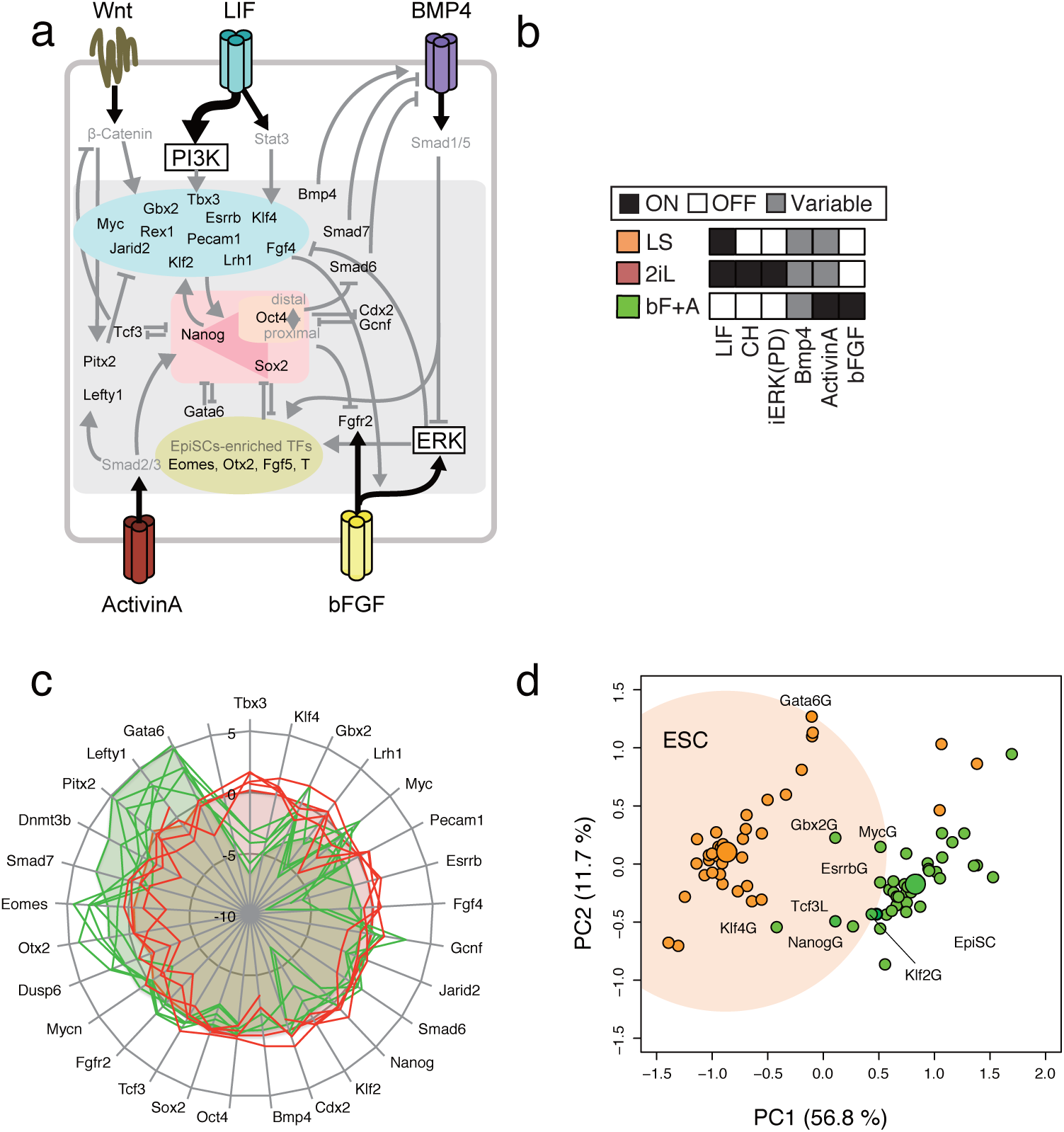
Simulation recapitulates distinct PSC states. **(a)** A schematic of the defined PSC gene/signal regulatory network model. The full representation of the model is shown in Supplementary Table S1 and Supplementary Notes Table M2. **(b)** Simulation inputs for LIF+Serum (LS: orange), 2i+LIF(2iL: red), and bFGF+Activin (bF+A: green) conditions. **(c)** Pinwheel diagram of relative population-averaged expression levels in predicted states (shaded area) under different input conditions (Red – mESC conditions, Green – EpiSC conditions) recapitulates experimental gene expression data (solid lines) from microarray and RNAseq studies. **(d)** *In silico* single gene GOF/LOF analysis of mESCs and EpiSCs was performed by fixing each gene in the GRN as ON or OFF, in either mESC (LS - orange) or EpiSC (bF+A - green) conditions. The calculated gene expression levels following each manipulation were mapped onto principle components. The individual gene perturbations that resulted in the changing of overall gene expression of EpiSCs to a more mESC-like one (green dots in orange shaded space) were predicted candidates for driving reversion from EpiSCs to mESCs.

Based on previous studies that reverse-engineered PSC-GRN to elucidate critical regulations ^33,34^, we performed refined Graphical Gaussian Modeling (GGM)^35^ to infer direct connectedness amongst genes using a collection of 1,295 publicly available microarray expression datasets for mESCs. (Supplementary Notes section 2). This resulted in a network of 29 genes including genes which are predicted to be highly correlated with the previously noted core pluripotency genes (Lefty1, Pitx2, Dusp6, Smad6 and Smad7) listed in Supplementary Table S1. Regulation directionality between network components was determined by either experimental evidence or categorizing based on gene functions (Supplementary Notes Table M1). Among 95 gene regulatory relationships identified through the GRN inference, the directionalities for ten remaining gene pairs left by previous experiments were determined by subsequent model selection based on fitting to reported single cell gene expression frequency. Taken together, reverse engineering-based GRN reconstruction, supplemented with manual curation led to an expanded GRN-signaling hybrid model consisting of 29 defined genes and 105 regulatory interactions between genes, seven signaling pathway activities and 24 regulations downstream of the signals (Supplementary Table S1 and Supplementary Notes Table M2).

Boolean logical functions of a target gene define the consequence of the binary states of its regulators with AND, OR, and NOT logic operators. Knowledge of all possible combinations and nesting of the operators significantly increase the number of possible models. Although this extends the capability of the model to describe a variety of regulatory topologies^19,21^, our focus was on applying a test model to the R-ABS/SCC approach to predict PSC GRN transitions in response to a wide range of signaling inputs. We used a biologically relevant and widely adapted rule where regulatory relationships were defined such that positive and negative inputs were combined using OR- and AND-functions, respectively. This rule states that a target gene will be present when one of its activators is present and concomitantly none of its repressors are present. Exceptions were made for genes whose coded proteins likely make a complex and work synergistically on regulating target gene expression (e.g. Oct4-Sox2 and Oct4-Cdx2). This resulted in 27,648 possible models including the unresolved directionalities of the above-mentioned 10 gene pairs, each of which has a distinct GRN topology. Among these possible models, we selected the top scoring model whose population-average gene expression level minimized Euclidean distance from single cell expression data of mESCs^1,36^ in standard culture conditions that contain LIF and fetal bovine serum (LS) (Supplementary Notes 3-5 and 3-6).

### Model recapitulates distinct PSC states

Using our GRN model and simulation strategy, we assessed the ability to predict PSC responses to different input signals. Mouse ESCs in LIF + serum medium (LS) were simulated by toggling LIF as continuously ON and allowing other endogenous signaling to undergo state transitions based on network structure (Fig 2b). Using the LS input rule, we identified only one SCC, which had no outgoing nodes. Five distinct steady-state attractors were also identified in LS. The predicted population-average gene expression levels of pluripotency-associated transcription factors were comparable with those reported using single cell RT-PCR in LS conditions (Fig S2a). Interestingly, the LS model also predicted that Oct4 was likely to co-exist with EpiTFs, while Sox2 showed strong negative correlation to EpiTFs, an observation consistent with previous reports^1,37^(Fig S2b).

To demonstrate the ability to predict alternate PSC states in response to changes in input signaling, we next performed simulations for EpiSCs ^16,38^ and naïve mESCs^39^ by changing only the input from LS to bFGF+Activin A (bF+A) or to LIF combined with inhibition of MEK and GSK3ß (2iL), respectively (Fig 2b). Simulations for both bF+A and 2iL conditions yielded only one PSC-associated SCC (see Methods and Supplementary Note section 5 for details). Notably, despite the fact that EpiSC gene expression data was not used to construct our generic PSC network, the model predicted distinct expression levels relative to those in mESC in LS, closely resembling experimental observations for the EpiSC state (Fig 2c, Fig S2c). Meanwhile, 2iL PSCs did not show significant differences in expression compared to LS PSCs, including expression of major pluripotency-supporting factors^40^, confirming that LIF is sufficient to maintain mESC-specific gene expression patterns. These data demonstrate that changing model inputs can drive GRN weighting to that observed in population-level *in vitro* experiments.

We next asked if we could manipulate GRN nodes directly and observe shifts between PSC states. This was done by setting individual genes ON (gain of function; GOF) or OFF (LOF), permanently, regardless of the states of their effectors. These simulations predicted Klf4, Nanog, Esrrb, Myc, and Gbx2 as drivers of EpiSC to ESC transition, and Tcf3 to be an inhibitor (Fig 2d, Fig S2d). These *de novo* results are consistent with previous experimental observations^14, 41–46^. The model also predicted that activating BMP4 while in bF+A conditions (i.e. EpiSC GRN) buoyed Oct4, Sox2, and Nanog (OSN) levels (Fig S2e); an observation that may explain the positive role of BMP4 in early stages of EpiSC reversion^41^. Taken together, we demonstrate the ability to quantitatively compare levels between different stable cell states by manipulating both extrinsic signals and/or endogenous GRN components.

### LIF stabilizes PSCs while 2i up-regulates OSN

Although LS and 2i with and without LIF (2iL and 2i-L, respectively) are sufficient to support stable PSCs^17,44,47,48^, and analysis of the GRN supported under these different input culture conditions are substantially similar^17,19^, clear morphological and phenotypic differences exist between cells cultured in baseline LIF + Serum vs. those supplemented with 2i (Fig 3a). We thus next sought to determine if our Boolean simulation approach could inform this biology using quantitative metrics related to GRN properties. To do this we developed mathematical formulations for three relevant metrics termed “*pluripotency*”, “*susceptibility*”, and “*sustainability*” (Supplementary Notes section 4). Briefly, *pluripotency* is defined as the population-average OSN expression level (sum of Oct4, Sox2, and Nanog levels). *Sustainability* is a score that reflects stability of an SCC in the absence of further perturbation and *susceptibility* quantifies the difference between an unperturbed SCC and an SCC with a perturbation of a GRN component (see “Calculation of population properties based on SCC” section in Methods for full formulations). These metrics serve as a foundation for quantitative comparisons of GRN properties, especially in dynamically stabilized cell states.

**Figure 3.**
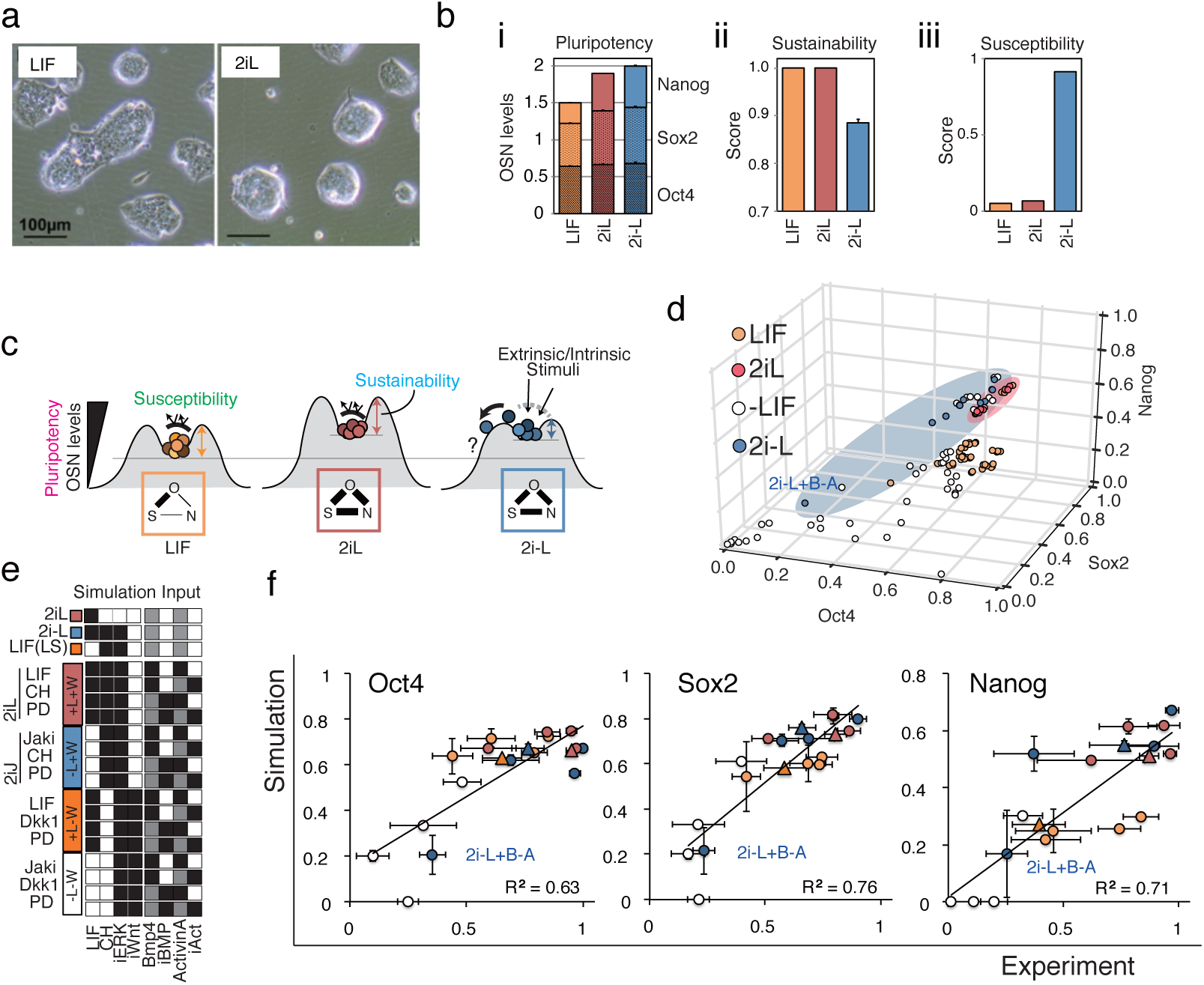
Dual inhibition (2i) supports the pluripotency core network (OSN) while LIF stabilizes PSCs. **(a)** Representative bright-field microscope images of mESC colonies in LIF and 2iL conditions with serum. **(b)** i. Pluripotency level (OSN expression) of each PSC-associated SCC. ii. Sustainability scores for each PSC-associated SCC. iii. Susceptibility of gene expression profiles against minimal perturbation to GRN topology was assessed *in silico* by measuring the change of variance in all genes. **(c)** Schematic illustration of the PSC metrics. The frequency of OSN-high cells reflects the population-level pluripotentiality. Sustainability reflects the intrinsic network stability in maintenance of the PSC state in the absence of extrinsic stimuli. Susceptibility measures the change of expression profiles to perturbations such as gene manipulations and signaling inputs and predicts the chance of PSC fate change. **(d)** Predicted population-averaged gene expression levels of OSN in SCCs from all possible combinations of signal inputs (White– without LIF, Orange – with LIF, Red – with 2iL and Blue – with 2i-L). **(e)** Four signaling pathways are manipulated in 16 conditions that are divided into four groups based on LIF- and Wnt-signal manipulations: +L+W (red, 2iL), –L+W (blue, 2iJ), +L–W (orange), and –L–W (white). Note that 2i+JAKi (2iJ) is the in vitro counterpart to the *in silico* 2i-L. **(f)** High content screening results of gene-expressing cell frequency (x-axis) and predicted population-average expression levels (y-axis) of OSN. Each condition is tested under activated and repressed Activin A/Nodal- and BMP-signals by addition of cytokines or inhibitors (±A±B). The symbol + indicates the addition of cytokines or small molecules that results in activation of the signaling pathway and the symbol – indicates the addition of inhibitors to pathway activity. Circles are 16 combinatorial signal conditions and triangles are the three control PSC conditions and are colorized using the same scheme outlined in (e). Experimental data is represented as mean and s.d. of three experiments, each performed in two replicates, and simulation data represents five independent simulations.

In the established stable PSCs supported by LS, 2iL and 2i-L, the *pluripotency* score was predicted to be higher in 2i(+/- L) than in LS (Fig 3b-i). This prediction was validated *in vitro* with immunocytochemistry for Oct4, Sox2, and Nanog (Fig S3a) and is consistent with data from previous reports^35^. The observed tighter correlations between pairs of individual OSN components in 2i-containing conditions, in which mESCs exhibited increased absolute levels and homogeneity of OSN than in the LS condition (Fig. S3b), indicate higher self-sustenance and homogeneity of the core network. Upon examining *sustainability,* our simulation scored the 2i-L model lower than the 2iL and LS models (Fig 3b-ii). As the sustainability score reflects the ability of a subpopulation in a given condition to maintain itself over time, the prediction infers that the presence of LIF raises the intrinsic stability of the subpopulation. This is consistent with the observations that LIF does not affect the pluripotency of mESCs in 2i-supplemented conditions, but enhances colony-forming efficiency^49,50^. Finally, to measure the *susceptibility* metric upon perturbations in GRN topology *in silico*, we tested the magnitude of expression changes by removing each single regulatory relationship from the original model network. This analysis identified key differences between 2iL and 2i-L. Overall, our simulation demonstrated that the GRN in 2i-L was more susceptible to perturbations of GRN topology than in conditions that contained LIF (Fig. 3b-iii and Fig.S3c). For example, the removal of the positive regulatory link from Nanog to Esrrb decreased OSN expression levels in 2i-L but not in LS and 2iL. This indicates that this link lacks built-in redundancy to be able to sustain OSN levels in the absence of LIF signaling. This confirmed the results from Dunn *et al.* that dual LOF of Nanog and Esrrb results in significant loss of pluripotency in 2i-L but not in 2iL^19^. Additionally, experimental validation using single and double gene LOF studies in 2i-L and 2iL performed in the study^19^ revealed that our model was able to accurately predict the outcomes of context-dependent LOF as well as the heightened susceptibility of mESCs cultured in 2i-L when compared to those cultured in 2iL (Fig S3d). Taken together, our simulations suggest that 2i drives PSCs into a naïve state expressing homogeneous levels of OSN, in part by supporting the OSN sub-network. Furthermore, the addition of LIF to 2i increased sustainability and decreased susceptibility of the overall GRN which, potentially by functional redundancy or additive effects^50^(Fig. 3c), are predicted to create barriers to the exit from pluripotency.

Based on our quantitative, metric-based analysis, we next hypothesized that pluripotent GRN supported by different input conditions (2iL and 2i-L) would be differentially susceptible to exogenous molecular perturbations. Indeed, simulation of all possible signal inputs (Fig. 3d) predicted that although 2i-containing conditions (2iL; red dots and 2i-L; blue dots) give overall higher OSN expression levels than +LIF (orange) or – LIF (black) conditions without 2i, the 2i-L condition has a higher variance of OSN levels (F=0.064, p-val= 8.0e-4; F=0.011, p-val= 2.2e-4; F=0.136, p-val= 1.1e-2 for Oct4, Sox2 and Nanog, respectively). Notably, OSN levels were predicted to decrease in the 2i-L condition only when combined with high BMP4 and low Activin A/Nodal (2i-L+B-A). To explicitly test these predictions, we measured core pluripotency GRN responses to combinations of four signaling inputs (LIF, BMP, WNT, and Activin A/Nodal)), both *in silico* (Fig.3e) and *in vitro* (Fig.S3e).To fully recapitulate simulation inputs, all signals that were turned OFF were validated with the corresponding small molecule inhibitor (e.g. -L in simulations = Janus kinase (JAK) inhibitor; JAKi (J) in experiments). Because LIF and WNT contribute to the maintenance of naïve mouse pluripotency^51,52^, we categorized each condition by the presence or absence of LIF and WNT signaling. Overall, OSN levels were high for conditions containing LIF or WNT, both in simulated and *in vitro* conditions (Fig S3f). Importantly, there was a high degree of correlation in OSN levels between *in silico* simulations and *in vitro* validations for all conditions tested (Fig 3f). While OSN levels were expectedly low in mESCs lacking both LIF and WNT signals and non-specific effects of the inhibitors may also make a contribution, the only condition robustly low in OSN in the presence of either LIF or WNT was the one that was additionally supplemented with the Activin signaling receptor (Activin receptor-like kinase 4/5/7, ALK4/5/7) inhibitor, ALKi, and BMP4 (2iJ+B-A). In conditions without LIF and with high WNT signaling (2iJ), OSN levels are sustained as high as the conditions with LIF and with low WNT signaling (Fig S3g). Interestingly, however, susceptibility of 2iJ conditions to perturbation by BMP and Activin signaling increased (Fig S3g, blue bars). These observations were conserved regardless of the presence of serum (Fig S3h). Overall these studies demonstrate that pluripotent cell states maintained under different input signaling conditions are differentially susceptible to pluripotency GRN destabilization, with the 2iJ conditions being particularly sensitive to perturbation.

### More permissive loss of pluripotency from 2i in the absence of LIF

We next set out to characterize the exit from pluripotency of PSC using the susceptible 2iJ-induced state. We first confirmed that mRNA levels of OSN are also decreased in 2iJ+B-A (2i-L+B-A *in silico*) with qRT-PCR after two days of culture in the respective conditions. Specifically, we found that the population-averaged gene expression levels of extra-embryonic lineage specifiers Cdx2 and Gata6 were significantly higher in the 2iJ+B-A condition than in the control conditions (2iL or 2iJ/2i-L) both in the simulation and the experiment (Fig 4a). Differentiation of naïve mESC to trophoblast stem (TS) cell–like cells occurs upon the forced expression of the trophoblast master regulator Cdx2, but not typically through the addition of media components^53,54^. However, apparent totipotency from mESCs derived in 2i^55^ has been reported, and SMAD1/5/8 activation helps drive trophoblast gene expression from mouse and human primed PSCs^38,56,57^. We thus asked if we could use our increased understanding of pluripotent cell state susceptibility to specifically direct the exit from pluripotency and access gene expression profiles normally reticent to differentiation from mESCs.

**Figure 4.**
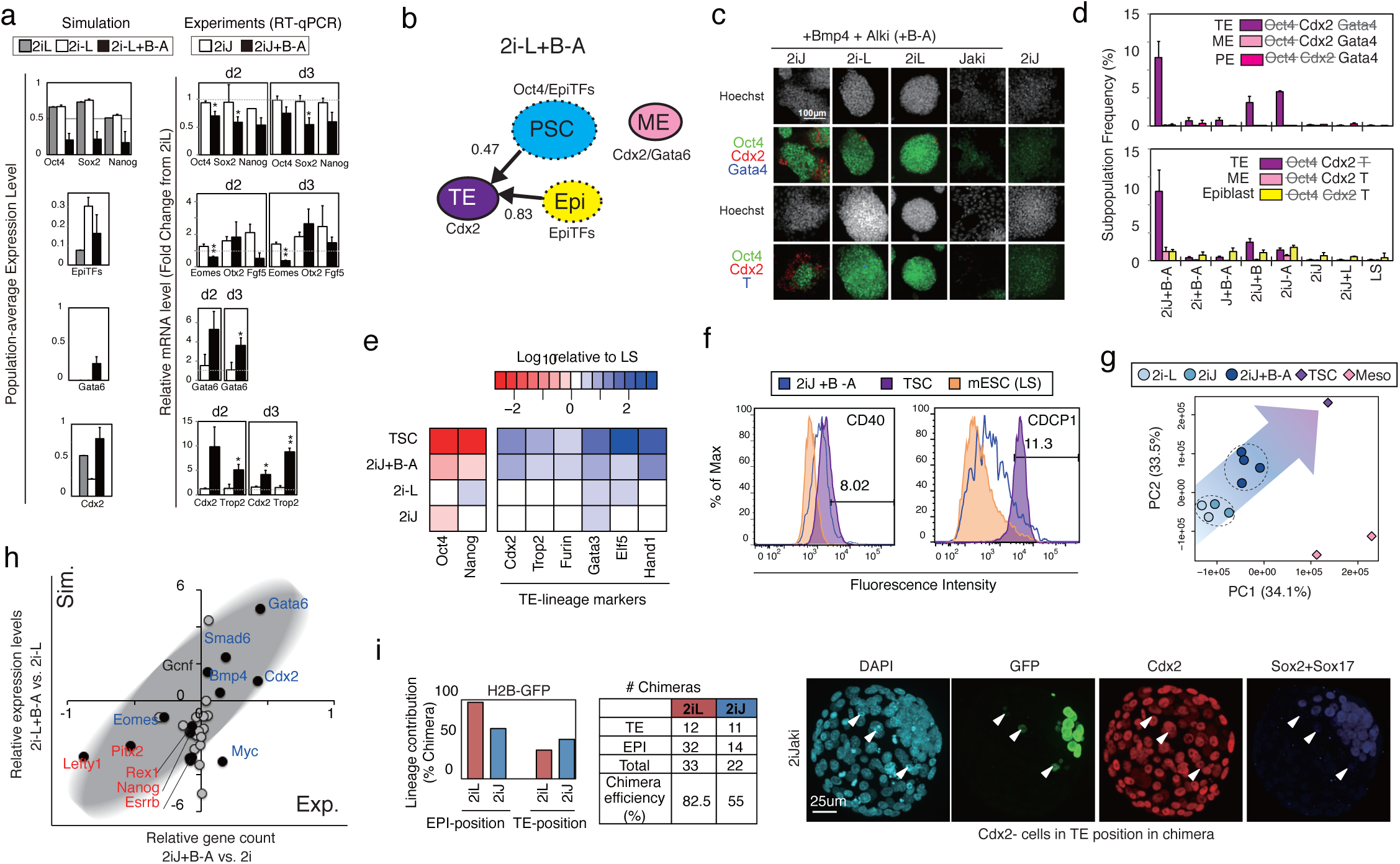
A culture-induced TE-like subpopulation. **(a)** Predicted population-level gene expression levels (left panel) and qRT-PCR-measured relative expressions (right panel; shown in fold-change from the levels of 2iL) of OSN and lineage markers. Data represents the mean and s.e.m. of three or four biological replicates; the differences between 2iJ and 2iJ+B were examined using a 2-tailed unpaired Student’s t test and asterisks indicate *p < 0.1 and ** *p* <0.05. **(b)** *In silico* subpopulation analysis via threshold-based characterization for individual SCCs under the input condition of 2i-L+B-A. Stable grouped profiles enriched as either PSC, TE, ME, PE or Epiblast-like subpopulations were traced in color-coded circles. The circles with solid line indicate SCCs with no outgoing edges (sustainability score =1.0), and those with dashed line indicate SCCs with lower sustainability. **(c)** Confocal images of immuno-staining of mESCs for Oct4/Cdx2/Gata4 (top) or Oct4/Cdx2/Brachyury (T) (bottom panels) cultured in the given conditions for 2 days. **(d)** Quantification of frequency of subpopulations that exhibit features of differentiation lineages. Data are means of three replicates and the results were confirmed in two independent studies. **(e)** qRT-PCR for pluripotency and extended TE-lineage maker genes. TSCs and mESCs after culture in each condition were compared. Data represents the mean of three replicates. **(f)** Flow cytometry histograms showing fluorescence intensity of CDCP1 and CD40 in individual samples of mESC in LS, TS, and mESCs cultured in 2iJ+B-A for 2 days. Percentage listed is that of positive cells in the 2iJ+B-A condition. **(g)** PCA plot of RNA-seq data for the top 40% of genes that show highest variance across all samples. Distinct cell types and conditions are indicated with different colors. Circles include day2 and day5 samples for 2i-L and 2iJ conditions, and two day 2 samples, day 5 and 11 for 2iJ+B-A condition. Diamonds indicate stable cell type no culture time defined. Meso indicates mesoderm progenitors. **(h)** Comparison of predicted gene expression levels and RNAseq-measured gene counts in 2iJ+B-A relative to 2iJ (Equivalent to 2i-L in simulation) for 29 genes involved in the model. The experimental mean relative gene expressions of day2 samples for the two conditions and the mean relative predicted levels are shown. Black dots indicate genes significantly up- or down-regulated (p < 0.05) in three 2iJ+B-A-treated samples compared with two 2iJ-treated samples. Genes in blue are up-regulated in TSC, and those in red are down-regulated in TSC compared with 2iJ-treated samples. **(i)** Left panel: *In vivo* lineage contribution frequency and chimera efficiency of H2B-GFP ESCs treated with either 2iL or 2iJ in the presence of serum. Lineage contribution efficiencies were calculated as number of chimeras with cells in Epiblast (EPI) or TE positions/total number of chimeras. Note that cells scored as “TE-position” did not express TE marker Cdx2. Chimera forming efficiency was calculated as number of chimeras/number of total aggregates made. Right panel: Representative images of aggregation chimeras at E4.5. We observed a number of cells in TE positions in chimeras. These cells ranged from live-looking to apoptotic, however none expressed Cdx2 and thus were not considered as viable, integrated contributions to the TE lineage.

Taking advantage of our framework’s capacity to predict the differentiation trajectories at exit from pluripotency, we scored individual SCCs and steady-state attractors that contained both highly expressed lineage markers and lowly expressed Oct4 as candidates for lineage bias. Cdx2 (Trophectoderm-associated – TE), Gata4/6 (Mesendoderm-associated – ME or Primitive Endoderm-associated – PE), and EpiTF genes (Post-implantation Epiblast) were specifically tracked (Fig 4b and Fig S4a).

To further explore conditions yielding induction of TE genes, we measured Cdx2 protein expression. As Cdx2 can emerge during primitive streak development^57^, we also co-stained with Oct4 and lineage markers Gata4 (endoderm) and Brachyury (mesoderm) (Fig 4c). There was a marked increase in the Cdx2 single-positive subpopulation in 2iJ+B-A, but not in the conditions lacking 2i, Jaki, BMP4, or ALKi (Fig 4d). The robust contribution of 2iJ+B-A condition towards a Cdx2-high state over time was confirmed by the frequency of Cdx2+/Oct4- cells after extending treatment to five days (Fig S4b). To further investigate this Cdx2+ state, we assayed a supplementary panel of TE markers by flow cytometry and by qRT-PCR. TE-enriched genes, such as Trop2 and bHLH transcription factor (Handl) also showed marked increase in expression in 2iJ+B-A relative to controls (Fig. 4e). TE-enriched surface markers CD40 and CUB domain-containing protein 1 (CDCP1)^58^ both increased in expression in 2iJ+B-A (Fig 4f and Fig S4c). Importantly, however, RNA-seq and subsequent principle components analysis (PCA) demonstrated that there exists a separation between mESCs in 2iJ+B-A and trophoblast stem cells (TSCs) (Fig 4g). This suggests that the fate transition of mESCs in 2iJ+B-A to a TE-like state is incomplete. Meanwhile, overall consistency between the simulation outputs and the differential expression profiles of mESCs in 2iJ+B-A from those in 2i were confirmed, as the significantly up-or down-regulated genes in 2iJ+B-A were almost universally predictable (Fig 4h).

Our analysis thus far demonstrates the strong predictive power of the simulation framework. We next set out to distinguish predictive gene expression changes from functional developmental states. Specifically we tested whether 2iJ-treated mESCs, which are in a pluripotent signal responsive state, would contribute to the developing embryo *in vivo* differently from other pluripotent conditions. We aggregated 2iJ-treated (2d) mESCs to totipotent host embryos (8-cell stage embryos) and allowed endogenous cues to guide differentiation of these cells during pre-implantation development. We noted an increased frequency of 2iJ-treated cells localizing to TE positions in the blastocyst compared to 2iL conditions (Fig 4i, left panel, Fig S4d), which was confirmed with different cell line^55^ (Fig S4), however these TE-positioned cells did not express Cdx2 at the time they were assayed in vivo and many seemed to have initiated apoptosis (Fig 4i, right panel). These results suggest 2iJ-treated cells are in an altered state of pluripotency, but are not fully competent to undergo trophoblast differentiation *in vivo,* either due to incomplete *in vitro* programing to a TSC like state or due to competing signals received *in vivo.* These differences between 2iJ/2iJ+B-A-treated mESCs and TSCs, as well as the contradictions between simulations and measurements suggest a requirement for TE-lineage specification on top of the Oct4-low/Cdx2-high state.

Taken together, the *in vitro* studies confirm the power of the model to predict the cell fate outcomes of PSCs exposed to complex exogenous signals. We demonstrate that the model can reveal new biology between different pluripotent cell states including EpiSC. We also demonstrate that mESCs cultured in 2i-L represent a unique cell state that exhibits high OSN levels (pluripotency) but are simultaneously highly responsive to a broad array of differentiation-inducing signals (susceptibility), and develop ad experimentally validate new metrics to distinguish between these states. Our system also can predict PSC differentiation transitions including into TE-like cells, as indicated by changes in gene and protein expression. Interestingly, our analysis also reveals conditions where additional signals or maturation steps may be required to fully transition cells across fate barriers, such as was observed for the TE-like cells.

## DISCUSSION

PSCs represent a powerful platform for the simulation of cell fate transitions^59^. A variety of methods have been used to model pluripotency. Prior models of mESC-GRNs with ordinary differential equations revealed mechanistic insights of cell fate transitions, but were limited in the number of network components around the core OSN network^60,61^. Recent work has provided an expanded number of network components in GRN models by use of model fitting and has derived network connectivity from either population-level knock-in/knock-down data^19^ or single cell expression data^21^. These models, however, were restricted in their ability to simulate transcriptional heterogeneity and exit from the pluripotent state. The absence of the interconnections between key signaling modalities for GRNs in these models can partly account for this shortcoming. To address this limitation, we used expression data from mESCs to populate our GRN framework and to produce a model of pluripotency that, when perturbed *in silico,* could recapitulate the emergence of subpopulations of cells observed under analogous *in vitro* and *in vivo* conditions. Additionally, by integrating principles of graph theory into our asynchronous Boolean model of pluripotency, we demonstrated the ability to model both exit from pluripotency and heterogeneity at two levels, the GRN level (fluctuation in gene expression) and the cell population composition level^62–64^. Importantly, the model with consensus interactions which excludes 10 predicted interactions but includes those validated from literature or ChIP-based genome interaction studies did not accurately predict the OSN levels in the various signal combinations in Fig.3f (Supplementary Notes 5-4). Also, deletion of any combination of two genes from the pluripotency supportive genes (Esrrb, Gbx2, Klf2 and Jarid2) in the model failed to predict the up-regulation of Cdx2 in the 2i-L+B-A condition. These results strongly support the validity of our PSC model as well as its ability to predict both known and novel effects of input signals.

The hierarchical differentiation process of PSCs is often illustrated by trajectories of cells in Waddington’s metaphorical landscape. Each basin is defined as an attractor, from which cells bifurcate into different downstream attractors that reflect differentiated cell types^65,66^. By quantifying *pluripotency, susceptibility,* and *sustainability* in our model, we propose that 2i conditions support a population of cells at the high potential (OSN) state. LIF signaling stabilizes cells within the stable local valley of the landscape by reinforcing the *pluripotency* GRN, increasing the threshold required to induce differentiation, and participating in the shaping of the landscape with regard to preferential (embryonic) versus non-preferential (extraembryonic) routes. Conversely, inhibition of JAK-STAT signaling destabilizes the *pluripotency* GRN and allows the expression of TE-lineage genes specifically in response to activation of BMP4 and inhibition of Activin A/Nodal signaling.

Although previous reports have demonstrated the ability of human and mouse primed pluripotent cells to differentiate into TE-like cells upon activation of BMP signaling^56,57^, these results may be condition and cell-line dependent and have not been connected to the underlying molecular structure of the pluripotency GRN. To date, there has been no evidence of the ability of cytokines and small molecules to drive TE differentiation from naïve pluripotent cells. A recent report from Morgani *et al.* demonstrated that mESC derived in 2i conditions, especially 2iL, increased the potential to contribute to extra-embryonic lineages including the TE-lineage *in vivo*^55^. Our results, however, showed greater ability to induce TE-lineage genes with active inhibition of JAK-STAT signaling in the presence of 2i, specifically with Bmp4 signal activation and Activin A/Nodal signal inhibition, a consequence of destabilization of the PSC GRN in a modified feedback signaling network environment. Our cells may require additional signals, such as Notch and Hippo^67^, and other TE differentiation signals that we did not use here (e.g. Elf5), to complete the programming. For example, DNA methylation of Elf5, a TE-specific transcription factor, silences its expression and blocks TE differentiation in mESCs, while the FGF-signal can initiate a positive feedback loop between CDX2 and Eomes through hypo-methylated Elf5 in TSCs^68^. Moreover, a higher epigenetic barrier may separate the TE-lineage and ESCs even in the hypomethylated ground state in 2i^69^. Nevertheless, our aim for the model was strictly to represent the exit from pluripotency in silico and validate the models predictions in vitro.

We anticipate that our simulation approach, which predicts changes in gene expression of sustained cell populations in response to signaling inputs, represents a broadly applicable approach to understanding the key control nodes triggering cell fate transitions. Beyond pluripotency, the use of our graph-theory based Boolean approach, together with combined modeling of signaling inputs and GRNs, may provide a new strategy for the prediction of aberrant cell fate transitions in normal or transformed somatic stem cells. In these systems, the exogenous influences typically described as components of the stem cell niche, serve to further broaden the likely regulatory feedback domain^70^. While the dynamic heterogeneity is conceptually supported by a series of analyses using single cell tracking techniques^7,23^, the empirical validation of the SCC approach, where the heterogeneous population of stem cells is assumed to correspond to SCC in the state transition graph of the asynchronously updated Boolean model, has not been performed as it requires live-cell tracking for multi-genes and multi-cells beyond our static cell profiling data collection. A recently reported in silico technique^73^, which derives GRN by retracing measured single cell expression profiles associated with asynchronous Boolean transitions, may possibly be used to evaluate how each SCC reflects inter-cell variability. However, these methods continue to face technical issues with respect to thresholding continuous gene expression and the quality of the single cell expression analysis itself. Another attractive approach is to extend the model into spatial-temporal metrics to validate predictions for self-organizing expression patterns in a system where dynamic stability of state-transitions derives inter-cell variability. As the framework is open to incorporating other known state stabilizing factors, such as epigenetic or metabolic control, it provides great opportunities to test hypothetical models of heterogeneity at both the genetic and cellular (i.e. tumor) level in cancer stem cells^70–72^.

## Acknowledgments

This work is funded by the Canadian Institutes of Health Research (CHIR grant #496640) (P.W.Z.). A.Y. has been supported by CHIR, G-COE and the JSPS institutional program for young researcher at Keio University, and Uehara Memorial Foundation, Japan. J.E.E.O. is supported by CHIR, IBBME international scholars program at University of Toronto, and Gålöstiftelsen, Sweden. E.P. is supported by RESTRACOMP Research Fellowship form Sickkids. P.W.Z. is supported as the Canada Research Chair in Stem Cell Bioengineering. We thank J. Chenoweth and R. McKay for RNAseq data on EpiSCs/mESCs, B. MacArthur, I. Lemischka, K. Hayashi, A. Surani for single cell data on mESCs, G. Martello and A. Smith for knockdown data on mESCs, C. Yoon for providing mesoderm progenitor cells, Dr. M. Nakanishi for initiating chimera analysis, and Dr. C. Bauwens for the critical reading and editing of this manuscript.

## Author Contributions

A.Y. designed and performed *in silico* study and bioinformatics analysis. K.O. designed and performed *in vitro* experiments. J.E.E.O. performed *in vitro* experiments and bioinformatics analysis. E.P. designed, E.P and M.N. performed, and J.R. supervised *in vivo* experiments. A.Y., K.O. and P.W.Z. designed the project and wrote the manuscript.

## METHODS

### Random Asynchronous Boolean Simulation (R-ABS)

Random ABS (R-ABS) has been performed using two assumptions: (i) every combination of model variables is equally likely to be calculated in a given step, and (ii) the state-space is generated with all the transition history of a sufficient number of consecutive steps from a sufficient number of random initial states. In this study, the random asynchronous simulation and the following calculation was done with 700 consecutive steps from 700 random initial states for one condition, where robust results can be derived from five independent simulations to reach similar population average expression probabilities (Supplementary Notes 5-3). Solving the Boolean equation in an asynchronous manner in each step is coded with Python using BooleanNet package ver.1.2.6 (http://code.google.com/p/booleannet/).

### Calculation of population properties based on strongly connected components (SCC)

Strongly connected components (SCC) are defined as clusters of unique expression profiles wherein all profiles are self-reachable and have more than one transitions to other profiles. Finding SCCs from the profile transition graph generated by R-ABS is done using networkx 1.9 python package.

For a particular SCC with *n* unique expression profiles, the transient matrix (*M*) with *n* rows and columns is defined. Each element (*m_ij_*) of *M* in row *i* and column *j* holds the value of the edge probability (i.e. accessibility from a source profile *j* to its target profile *i*) ranging from 0 to 1, which represents the relative transition frequencies from a specific expression profile to one of its target profiles among all transitions from the source profile. The profile-probability *v_i_,* of profile *i* indicates the chance that a certain cell resides at profile *i* in the SCC. The sum of the products of *m_ij_* and *v_j_* indicates the profile-probability (*V_i_*) of the source profile *j*, which has a transition path to *i*. The distribution approaches a limiting distribution *v*, where *v* =*M* × *v* is satisfied. Assuming the cell population is a sum of probabilities of heterogeneous single cell states (profiles), Σ_*n*_(*v_n_*)is equal to 1. Solving v under these constraints gives the principal eigenvector of *M* that tells us which profile is likely to be arrived at after a certain number of simulation steps from any profile in the SCC.

Sustainability for a particular SCC indicates the probability of out-going cells from the dynamic stable state, reflecting a quantitative measure of the intrinsic stability of the GRN within the SCC. The sustainability score (*S_scc_*) is defined as *S_scc_ =* 1 - (Σ_*j*_(*ν_j_*) × Σ_*k*_ν_*jk*_)), ranging from 0 to 1, assuming profile *j* (inside the SCC) has out-going edges towards profile *k* (outside the SCC). The expression frequency (*p*) of model component (g) in a particular SCC can be calculated as a summation of all v with ON-expression of the gene:

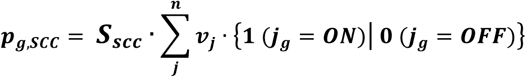

where *j_g_* denotes the binary state of *g* in the profile *j*. To avoid overestimation of *v* and to maintain calculation accuracy of population-average expression level, we defined the thresholds for SCCs to be considered in the analysis as the number of profiles >10 and sustainability score > 0.7. As a larger dynamic stable state of PSCs is more likely to exist over time it will become a larger determinant of population-average expression levels. Consequently, population-level gene expression level is calculated by averaging multiple SCCs:

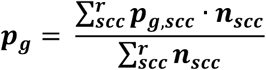

where *r* is the number of SCCs found under the given condition satisfying the predefined threshold, and *n* is the number of unique profiles involved in the SCC. A small constant value for *p* (*p*=1e-5) was applied where necessary to avoid zero division.

The subpopulation characteristics were quantified based on the SCC-averaged expression levels of lineage and pluripotency associated genes (Cdx2, EpiTFs, Gata6 and Oct4) in the associated SCC. Each SCC and steady-state attractor is classified based on the thresholds on the SCC-averaged expression levels (Cdx2: 0.7, EpiTFs: 0.2, Gata6: 0.5 and Oct4:0.3). High expression of individual markers in separate SCCs is likely to represent TE (Cdx2), Epiblast (EpiTFs), PE (Gata6) and PSC (Oct4), while high co-expression of differentiation lineage markers (Cdx2, EpiTFs, Gata6) in the same SCC would suggest a ME lineage (Morgani, 2013). The population-averaged subpopulation frequency is calculated by simply applying the same strategy with the calculation of expression level for each SCC where the size of the SCC (the number of unique expression profiles) and the sustainability score are taken into consideration:

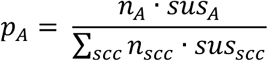

where the number of unique profiles in the SCC *A* and sustainability of *A* are indicated as *n_A_* and *sus_A_,* respectively.

### GRN inference

We collected mESC expression data on 1,295 genes using the Affymetrix Mouse 430 2.0 Array from the Gene Expression Omnibus (GEO) database at the US National Center for Biotechnology Information (NCBI) and ArrayExpress at the European Bioinformatics Institute (EBI) (See Appendix of the Supplementary Note). Graphical Gaussian Modeling (GGM) was employed to infer direct regulatory networks between gene pairs based on partial correlations. All data including 45,101 probe sets were quantile normalized with the R/Bioconductor limma package. The probe sets were then converted into 13,879 unique genes by taking mean values of probes annotated as same gene. Twenty thousand iterations were performed where 1,000 genes were randomly sampled for one partial correlation analysis with the GeneNet package in R. The lowest partial correlation for a certain gene pair among whole iterations satisfying constraints (Pearson’s correlations > |0.3|, *p* < 0.05) was considered as a GGM score. The gene-to-gene links with positive/negative-high GGM scores (> 0.03) were taken into consideration as regulatory edges in the model.

### Model selection

The candidate models were evaluated by comparing the Euclidean distance between predicted population-average expression levels and observed frequency of gene-expressing single cells in single-cell data sets^1,36^. Both the simulation and the experiment were performed in the control LS condition. The mRNA expression of single cells were binarized by applying k-means clustering on the expression data across all samples with k=2, and then the frequency of gene-expressing single-cells in the population were calculated. For the k-means clustering, we used the cluster.vq function in the Python scipy package. As there are two series of single-cell transcriptomic reference profiles, the average values of the frequencies from the two experiments’ datasets were used.

### Code availability

Python codes used for the simulation and SCC analysis of population properties are available upon request.

### Cell Culture

R1 mouse embryonic stem cells (mESCs) were cultured in serum-containing and feeder-free conditions as described previously^74^. Validation of predicted responses to exogenous signaling was performed in serum-containing medium supplemented with combinations of the following cytokines/small molecules: LIF (Millipore ESG1107 – 10ng/ml), JAK inhibitor (EMD Millipore 420097 – 2.0μM), BMP4 (R&D Systems 314-BP-010 – 10ng/ml), LDN193189 (Reagents Direct 36-F52 – 0.1μM), CHIR99021 (Reagents Direct 27-H76 – 3μM), Dkk1 (R&D Systems 1765-DK-010 – 275ng/ml), bFGF (Peprotech 100-18B -20ng/ml), PD0325901 (Reagents Direct 39-C68 – 1μM), Activin A (R&D Systems 338-AC-050 – 20ng/ml), and ALK5 inhibitor II (Enzo Life Sciences ALX-270-445 – 10μM and Cedarlane ALX-270-445 for RNA-seq). Trophoblast stem cells (TSCs) were cultured as described previously^75^. Mesoderm progenitor cells were generated from embryoid bodies (EBs) in differentiation medium containing Iscove’s modified Dulbecco’s medium (IMDM; Thermo Fisher Scientific) and Ham’s F-12 nutrient mix (Thermo Fisher Scientific) supplemented with 1X B-27 supplement (Thermo Fisher Scientific), 1X N-2 supplement (Thermo Fisher Scientific), 2 mM Glutamax (Thermo Fisher Scientific), 100 U/mL penicillin-streptomycin (Thermo Fisher Scientific), 0.05% bovine serum albumin (BSA; Wisent), 150μM monothioglycerol (MTG; Sigma), and 0.5 mM ascorbic acid (Sigma). On day 2, EBs were harvested, dissociated into single cells, and re-aggregated in 100 mm Petri dishes (BD Biosciences) with differentiation medium further supplemented with BMP4 (1 ng/mL), Activin A (2 ng/mL), and Wnt3a (3 ng/mL). Mesoderm progenitors were isolated either before or 24 hrs after addition of IWP-2 (Reagents Direct 57-G89 - 2μM) to each Petri dish on day 3.

### mRNA Quantification with qRT-PCR

Primer sequences were obtained from PrimerBank^76^ and are listed in Supplementary Table S2. qRT-PCR was carried out as described previously^15^. Briefly, cells were lysed and RNA was isolated using the PureLink^®^ RNA Mini Kit (Life Technologies). RNA was converted to cDNA using SuperScript III Reverse Transcriptase (Life Technologies) and amplified in FastStart SYBR Green Master Mix (Roche) using the 7900HT Fast Real-Time PCR System (Thermo Fisher) with an annealing temperature of 60°C. Each data set was normalized to ß-actin in each condition and then normalized to the control.

### *In vitro* Immunostaining and Quantification

Cells were fixed and stained as described previously^15^. The following antibodies were used at a 1:200 dilution: Oct4 (BD biosciences 611203), Oct4 rbIgG1 (Cell Signaling 2840S), Sox2 (R&D Systems MAB2018), Nanog (eBiosciences 14-5761-80), Cdx2 (BioGenex MU392-UC), Gata4 (Santa Cruz Biotechnology sc-1237), and Brachyrury (R&D Systems AF2085). Stained cells were quantified on the Cellomics™ (ThermoFisher) high content screening platform.

### Flow Cytometry

Cells (mESCs and TSCs) were first stained for surface markers CDCP1 (R&D Systems AF4515) and CD40 (BD Biosciences 562846) using antibodies at 1:100 dilutions and assayed using flow cytometry (BD LSRFortessa). Cells were also stained with a live/dead stain (LIVE/DEAD^®^ Fixable Far Red Dead Cell Stain Kit for fixed cells, 7-AAD for live cells – Life Technologies) and gated for live cells. Final graphs were generated using FlowJo software.

### RNAseq

RNA was extracted using the PureLink RNA Mini Kit (Ambion, Life Technologies, Cat no. 12183018A and 12183025) according to the manufacturer’s instructions. Cells were homogenized using a QIAshredder (Qiagen, Cat no. 79654). Cell pellets were frozen at the treatment-specific time points and RNA was extracted from all pellets at the same time for each analysis. Quality control of total RNA was done on an Agilent Bioanalyzer 2100 RNA Nano chip following Agilent Technologies’ recommendation. RNA libraries were then sequenced on an Illumina HiSeq 2500 platform using a High Throughput Run Mode flowcell and the V4 sequencing chemistry following Illumina’s recommended protocol to generate paired-end reads of 126-bases in length. Reads were trimmed for adapters and a phred33 quality cutoff of 20 using TrimeGalore with cutadapt, and mapped to the Ensembl NCBIM37 mouse genome using STAR 2.4.2a. To adjust batch effects between experiments of two distinct days, COMBAT, an R package for Empirical Bayes method (http://statistics.byu.edu/johnson/ComBat/) was utilized.

### Chimera Generation and Analysis

E14 Ju09 HV H2B-Tomato ESCs^55^ were a gift from Joshua Brickmann and CAG H2B-eGFP ESCs were derived from mice^77^. ESCs were maintained in ESC medium containing 2i/LIF/serum. Cells were passaged twice with 2iL on mouse embryoic fibrobalasts and inactivated with no growth factor medium. Cells were then treated for 48 hours with 2i/LIF/serum or 2i/Jaki/serum on mouse embryonic fibroblasts. Chimeric embryos were generated by morula aggregation. Clusters of 5 to 10 ESCs were aggregated with wild-type CD1 morulae and cultured in Potassium (K) Simplex optimized media (KSOM; Chemicon) under paraffin oil at 37°C and 5% CO2 until the late blastocyst stage (embryonic day 4.5). Blastocyst embryos were subjected to immunofluorescent staining using anti-Cdx2 (1:600, Abcam ab76541), anti-Gata4 (1:100, Santa Cruz Biotech sc- 9053), anti-Sox2 (1:100, R&D Systems AF2018), anti-GFP (1:400, Abcam ab13970) and anti-RFP (1:100, Abcam ab65856) antibodies. Imaging was performed using a Quorum WaveFX spinning disk confocal system and Volocity acquisition software (Perkin Elmer). The frequency of cells localizing to extra-embryonic - trophectoderm (TE) positions in the blastocyst was counted. The investigators were not blinded to allocation during outcome assessment, and the experiments were not randomized.

### Data and Statistical Analysis

We assume that each well of a culture dish behaves as a biological replicate. No statistical methods were used to predetermine sample sizes. Images including immuno-staining experiments shown are representative of at least three independent experiments. Simulation data were derived from five individual runs for the indicated inputs. For the calculation of *p*-value, Wilcoxon exact rank test (R: coin package) was used for comparison of data groups unless otherwise stated. All tests of statistical significance were two-sided (* p <.05, ** p<.01).

### Data Availability

RNA-seq data have been deposited in the Gene Expression Omnibus (GEO) under the accession number of [TBD].

## Supplementary Figures

**Supplementary Figure S1.**
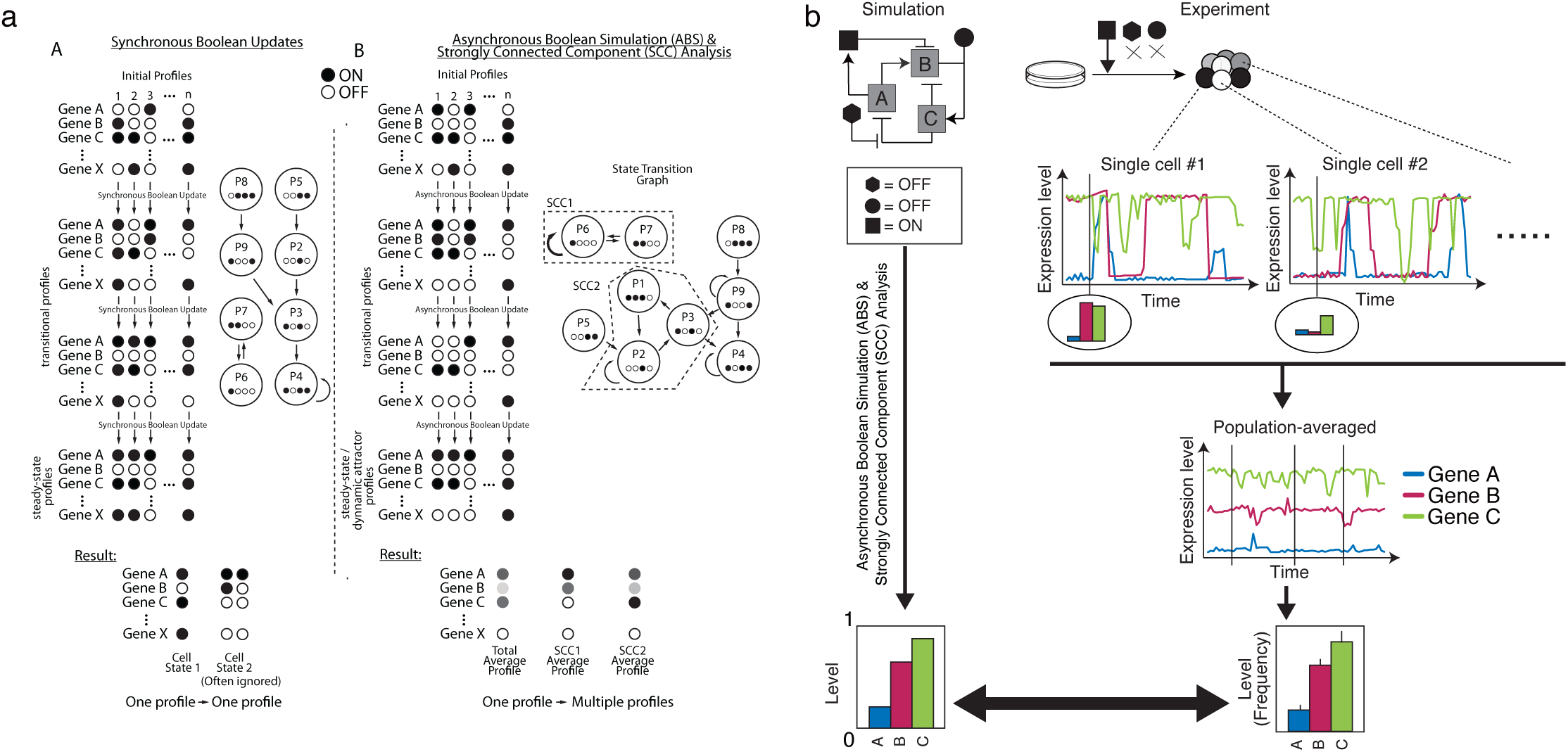
General strategy of the simulation. **(a)** The property of synchronous (A) and asynchronous (B) Boolean simulation. Importantly in ABS, a single starting profile can give rise to multiple different resultant profiles and therefore all transitional profiles are represented with probabilities. Note that we used random ABS where the genes updated in each transition are set randomly to reduce the calculation cost. **(b)** The background of our simulation framework: heterogeneity derived from single cell fluctuations and stabilization of PSC populations showing robust gene expression pattern under a certain signaling input condition (depicted with black geometric shapes).

**Supplementary Figure S2.**
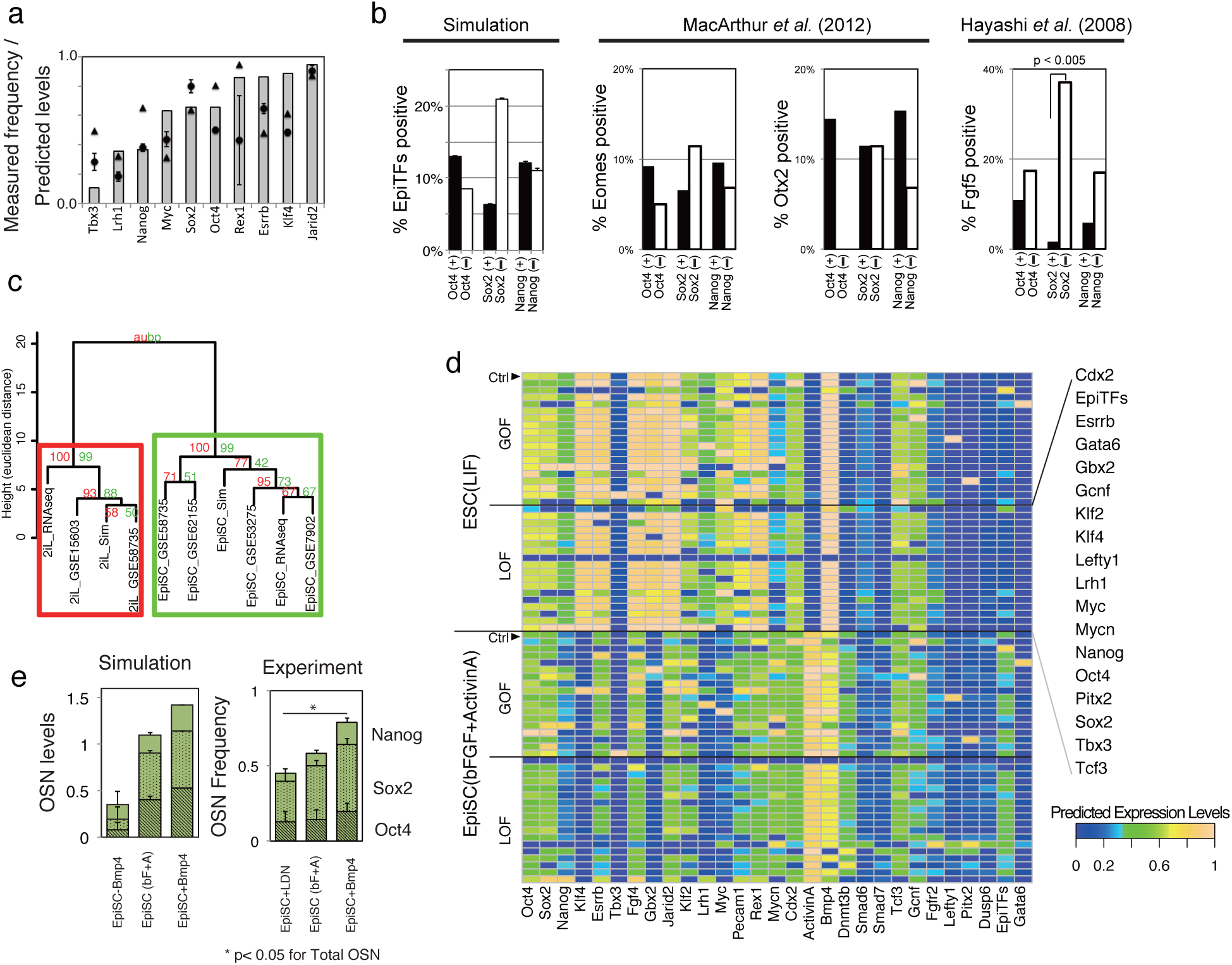
Comparison of predicted and experimentally observed data on gene expression patterns in distinct PSCs; related to Figure 2. **(a)** Predicted population-average expression level (mean of five independent simulations) for each pluripotency-associated gene in the control LS condition is comparable with the frequency of gene-expressing cells from the reported single cell measurements using RNAseq (triangle)^36^ and fludigm-qPCR^10^(circle)^1^. The gene expression levels were binarized into ON or OFF for each single cell to calculate frequencies. **(b)** Comparison of predicted co-expressions of epiblast-specific genes (EpiTFs – Eomes, Otx2, and Fgf5) with O/S/N and those measured by qPCR-based single cell mRNA data. Oct4 and Nanog are likely to be co-expressed with EpiTFs whereas Sox2 is not, as it shows a negative correlation with EpiTFs (i.e. EpiTF expression in Sox2- cells is higher) in both our simulation and in published, single cell expression data^1,37^. *p*-value was calculated by Fisher's exact test. **(c)** Simulation of distinct PSCs recapitulates population-level gene expression measurements. All expression data is scaled relative to respective gene expression in control LS conditions. The hierarchical clustering with AU (Approximately Unbiased) *p*-value and BP (Bootstrap Probability) value based on multiscale bootstrap resampling was carried out with the pvclust package in R. The results are displayed in the pinwheel view in **Fig 2c**, where upper and lower limits are set to +5 and −10, respectively, in the simulation to avoid singularities caused by null expression. The experimental data was taken from indicated GEO entries and RNAseq data for EpiSCs was kindly provided by Dr. Ronald Mckay. Cell lines used are 129 (GSE15603 and GSE62155), J1 (GSE58735), ESF175/1,ESF58/2, ESF122 (GSE7902) cell lines, or mESCs were derived from C57BL/6J strain blastocysts (GSE53275). **(d)** *In silico* GOF/LOF study in EpiSC (bF+A) or mESC (LS) conditions. All SCCs above ten profiles and sustainability > 0.7 are shown, and the gene expression levels of each component in each SCC are color-coded between blue (0.0) to yellow (1.0). The population-average expression levels based on the GOF/LOF results were shown in the PCA-metrics in Fig.2c. **(e)** Predictions (left) and measurements (right) of population-averaged expression levels of OSN in EpiSC-conditions (bF+A) increased in response to extrinsic manipulation of BMP4. BMP4 was set as continuous-ON (EpiSC+BMP4) and random (EpiSC). The frequencies reported for Oct4, Sox2 and Nanog-positive cells, assessed by Cellomics high content screening, represent the mean and s.d. of four replicates. Asterisk indicates the significant difference (p < 0.05 unpaired, two-sided t test for total OSN).

**Supplementary Figure S3.**
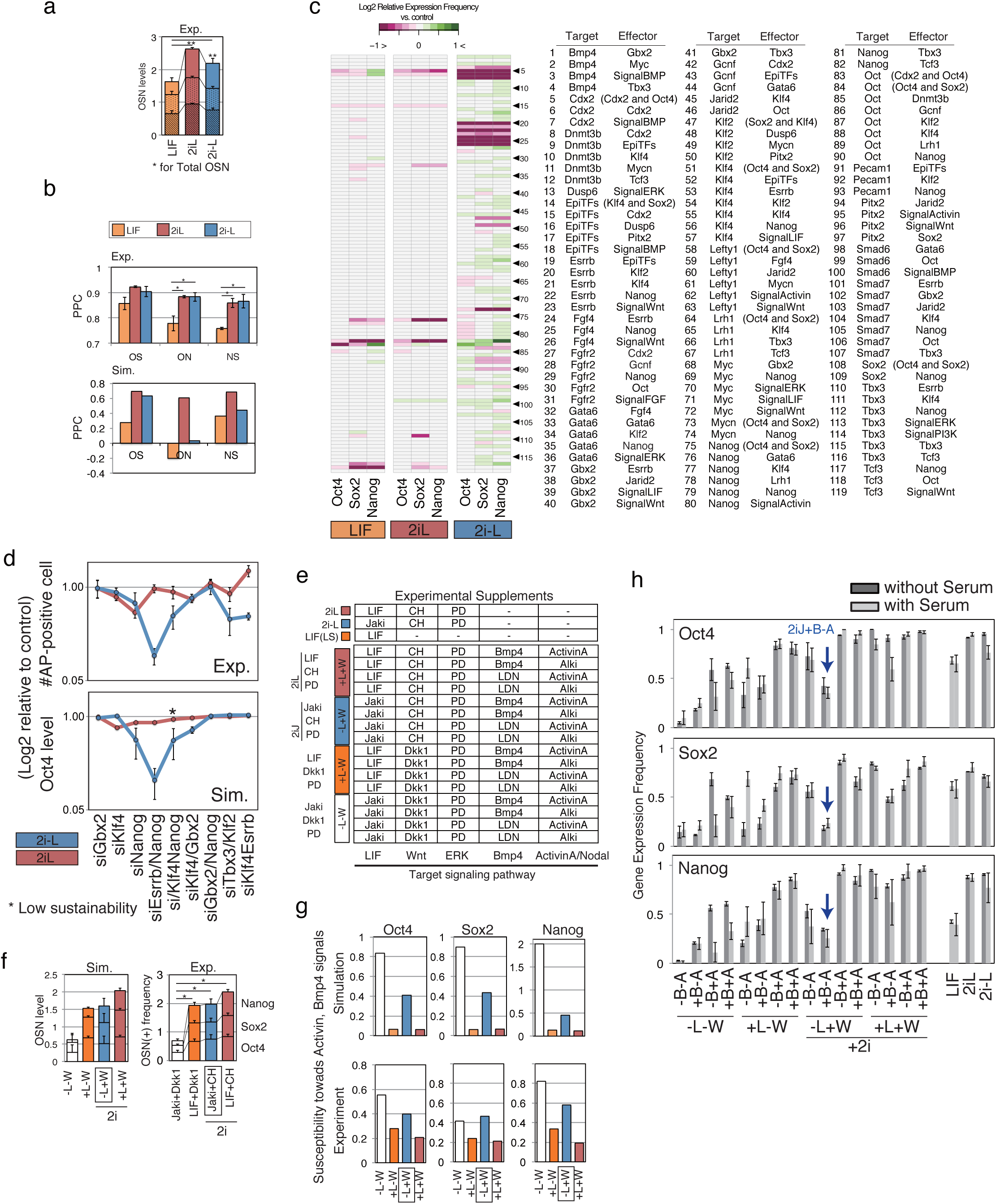
LIF stabilizes the pluripotent population while 2i up-regulates OSN; related to Figure 3. **(a)** Summation of frequencies (0-1) of O/S/N-positive cells assessed by high content screening at 2 days after medium replacement (n=6). Asterisk indicates the significant difference for total OSN. **(b)** Upper panel: A measure of Pearson’s correlation for OSN in 2i-supplemented conditions, quantified using high content screening (n=6). Lower panel: A measure of Pearson’s correlation for OSN in the results of a minimal perturbation-sensitivity analysis described in c. **(c)** The results of minimal perturbation-sensitivity analysis of the model where the model network was perturbed by removing single regulatory edges. Purple indicates down-regulation of the genes by removing the regulatory edge, which means the regulation had a positive role on the expression level of O/S/N, while green indicates the reverse. Note that the results shown are the effects of regulatory edge-removal: the removal of inhibitory regulation from Cdx2 to Oct4 increases Oct4, which means that the regulation edge acts as a negative effector for Oct4 level. **(d)** The difference in PSC population stability between 2iL and 2i-L conditions was assessed via published single gene-LOF and double genes-LOF *in vitro*^19^ and *in silico.* The upper panel depicts the experimental results for the relative count of alkaline phosphatase (AP) positive cells to untreated colonies upon gene manipulations. The simulation data (lower panel) shows population-average expression level of Oct4 relative to controls (2iL and 2i-L) where the manipulated gene was set as continuously OFF. Blue nodes indicate the response in 2i-L while red nodes indicate those in 2iL (n=5 for simulation and n=3-5 for experimental). **(e)** The experimental inputs (additives of the conditions) corresponding to the simulation inputs shown in **Fig 3e**. Abbreviations used are as follows: Jaki for Janus kinase (JAK) inhibitor, Dkk1 for Dickkopf1 and Alki for chemical inhibitor selective for activin receptor-like kinase (ALK) 4/5/7. **(f)** Predicted levels (left) and measured gene-expressing cell frequencies (right) of O/S/N for each condition group (L = LIF; W = WNT). The average value of four distinct signal conditions (±BMP±Activin/Nodal) in each group for each gene was calculated and summed up into an OSN score (n = 6). Asterisk indicates the significant difference for total OSN. **(g)** Susceptibility of O/S/N expression levels of PSCs to Activin and BMP signal perturbations was predicted (upper panel) and measured (lower panel). Standard deviation per mean of predicted expression levels or gene-expressing frequencies from immuno-staining was calculated as a coefficient of variation for each group (±LIF±WNT) including four distinct signal conditions (±BMP±Activin/Nodal). **(h)** Frequencies of O/S/N-expressing cells in the 19 combinatorial signal conditions with and without serum assessed by immuno-staining (n=2). Most conditions were equivalent except for –L–W– B+A and +L–W–B–A. The notable down-regulation of OSN in –L+W+B–A (=2iJ+B–A) was seen in both with- and without-serum conditions.

**Supplementary Figure S4.**
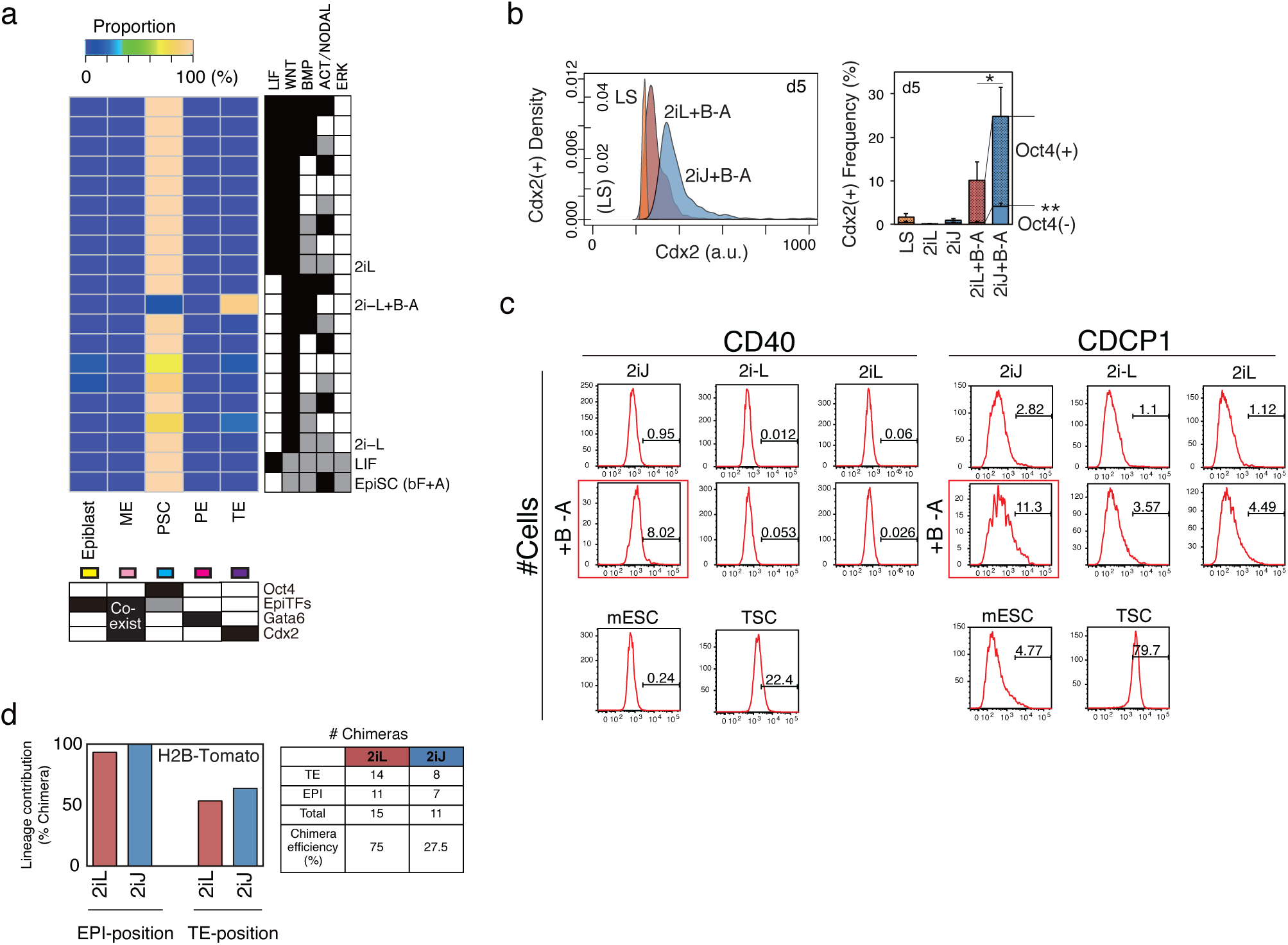
*In silico* subpopulation analysis reveals mESCs exit pluripotency towards TE-like lineage in 2iJ+B-A condition; related to Figure 4. **(a)** *In silico* subpopulation analysis of possible signaling input combinations with 2iL and 2i-L. The threshold values for predicted expression levels of lineage specifiers in each SCC are set as follows: Oct4=0.3, EpiTFs=0.2, and Gata6=0.5 and Cdx2=0.7. **(b)** Frequency of Cdx2+ population including TE-like sub-population (Cdx2+/Oct4-) after five days in culture measured by immuno-staining. Data represents the means of six replicates consisting of two biological replicates with three technical replicates in each. The representative density plot of immuno-staining of Cdx2 on day 5 is shown in the left panel. **(c)** Histograms for expression of TE-enriched cell surface markers, CD40 and CDCP1. Shown are mESCs treated in 2iJ (1^st^ column), 2i-L (2^nd^ column), or 2iL(3^rd^ column), in unsupplemented medium (1^st^ row) or BMP4 and ALKi supplemented medium (+B-A) (2^nd^ row). Control mESCs (kept in control LS) and TSCs are shown at the bottom. **(d)** *In vivo* lineage contribution frequency and chimera efficiency of H2B-Tomato ESCs^55^ treated with either 2iL or 2iJ in the presence of serum.

**Supplementary Table S1:**
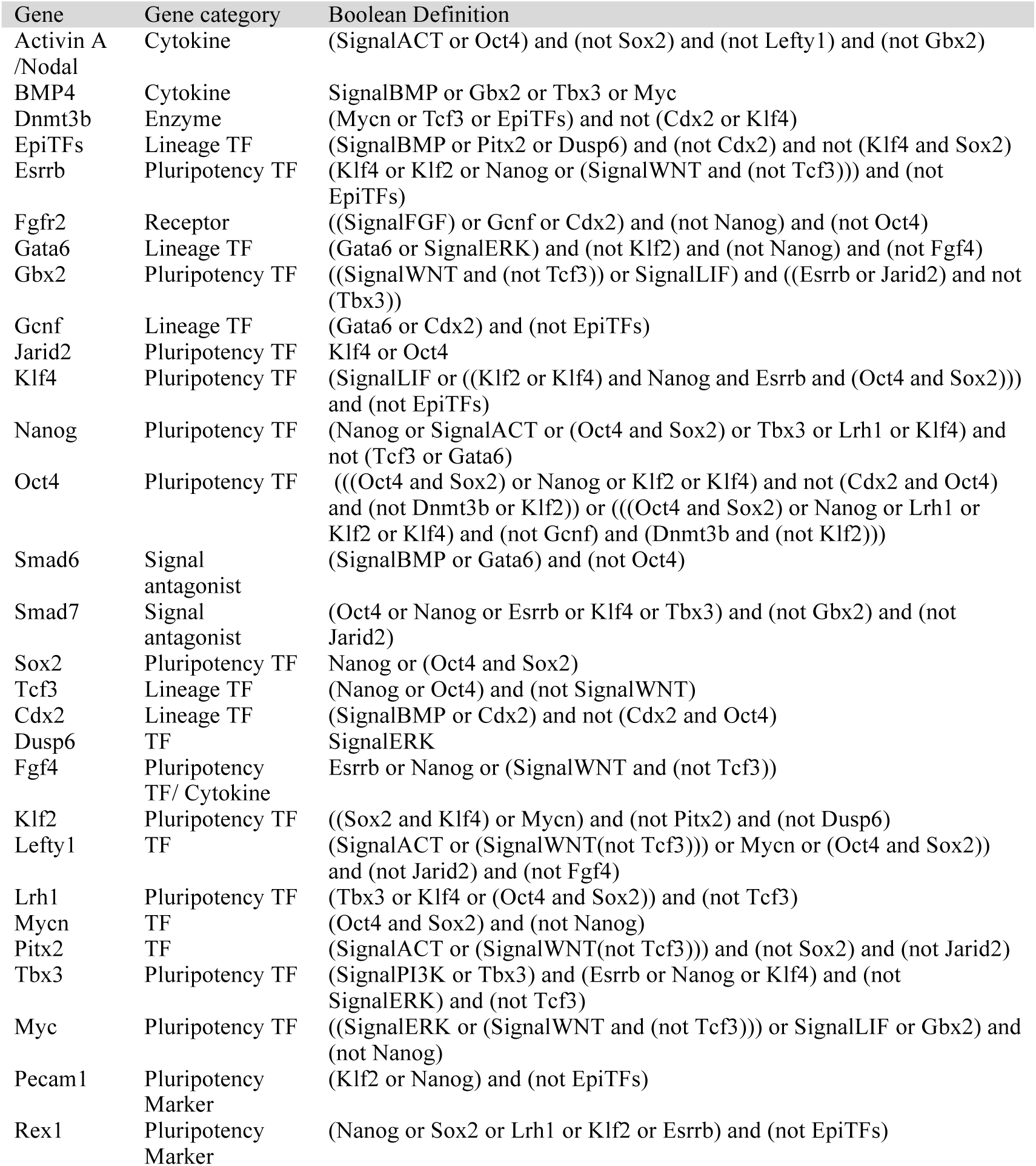
Boolean definition for genes in the model.

**Supplementary S2:**
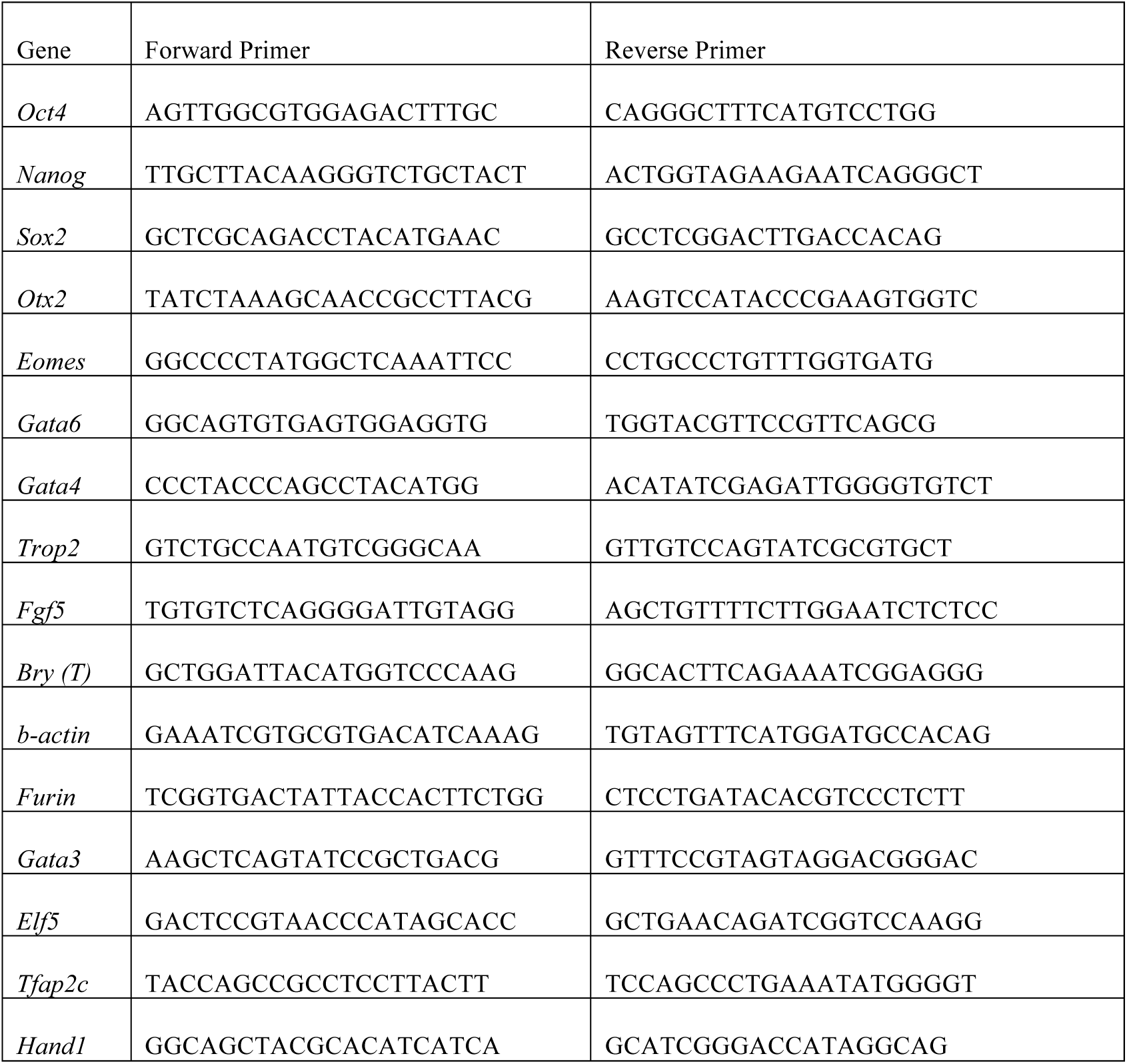
Primer sequences used in the study.

## References

1. MacArthur, B. D. et al. Nanog-dependent feedback loops regulate murine embryonic stem cell heterogeneity. Nat. Cell Biol. 14, 1139–1147 (2012).

2. Gupta, P. B. et al. Stochastic state transitions give rise to phenotypic equilibrium in populations of cancer cells. Cell 146, 633–644 (2011).

3. Hoppe, P. S., Coutu, D. L. & Schroeder, T. Single-cell technologies sharpen up mammalian stem cell research. Nat. Cell Biol. 16, 919–927 (2014).

4. Miyanari, Y. & Torres-Padilla, M.-E. Control of ground-state pluripotency by allelic regulation of Nanog. Nature 483, 470–473 (2012).

5. Schroeder, T. Long-term single-cell imaging of mammalian stem cells. Nat. Methods 8, S30-35 (2011).

6. Eldar, A. & Elowitz, M. B. Functional roles for noise in genetic circuits. Nature 467, 167–173 (2010).

7. Singer, Z. S. et al. Dynamic heterogeneity and DNA methylation in embryonic stem cells. Mol. Cell 55, 319–331 (2014).

8. Rompolas, P., Mesa, K. R. & Greco, V. Spatial organization within a niche as a determinant of stem-cell fate. Nature 502, 513-518 (2013).

9. Chambers, I. et al. Nanog safeguards pluripotency and mediates germline development. Nature 450, 1230–1234 (2007).

10. Toyooka, Y., Shimosato, D., Murakami, K., Takahashi, K. & Niwa, H. Identification and characterization of subpopulations in undifferentiated ES cell culture. Dev. Camb. Engl. 135, 909–918 (2008).

11. Ng, H.-H. & Surani, M. A. The transcriptional and signalling networks of pluripotency. Nat. Cell Biol. 13, 490–496 (2011).

12. Moledina, F. et al. Predictive microfluidic control of regulatory ligand trajectories in individual pluripotent cells. Proc. Natl. Acad. Sci. U. S. A. 109, 3264–3269 (2012).

13. Davey, R. E. & Zandstra, P. W. Spatial organization of embryonic stem cell responsiveness to autocrine gp130 ligands reveals an autoregulatory stem cell niche. Stem Cells Dayt. Ohio 24, 2538–2548 (2006).

14. Guo, G. et al. Klf4 reverts developmentally programmed restriction of ground state pluripotency. Dev. Camb. Engl. 136, 1063–1069 (2009).

15. Onishi, K., Tonge, P. D., Nagy, A. & Zandstra, P. W. Microenvironment-mediated reversion of epiblast stem cells by reactivation of repressed JAK-STAT signaling. Integr. Biol. Quant. Biosci. Nano Macro 4, 1367–1376 (2012).

16. Tesar, P. J. et al. New cell lines from mouse epiblast share defining features with human embryonic stem cells. Nature 448, 196–199 (2007).

17. Nichols, J., Silva, J., Roode, M. & Smith, A. Suppression of Erk signalling promotes ground state pluripotency in the mouse embryo. Dev. Camb. Engl. 136, 3215–3222 (2009).

18. Evans, M. Discovering pluripotency: 30 years of mouse embryonic stem cells. Nat. Rev. Mol. Cell Biol. 12, 680–686 (2011).

19. Dunn, S.-J., Martello, G., Yordanov, B., Emmott, S. & Smith, A. G. Defining an essential transcription factor program for naïve pluripotency. Science 344, 1156–1160 (2014).

20. Okawa, S. & del Sol, A. A computational strategy for predicting lineage specifiers in stem cell subpopulations. Stem Cell Res. 15, 427–434 (2015).

21. Xu, H., Ang, Y.-S., Sevilla, A., Lemischka, I. R. & Ma’ayan, A. Construction and validation of a regulatory network for pluripotency and self-renewal of mouse embryonic stem cells. PLoS Comput. Biol. 10, e1003777 (2014).

22. Kalmar, T. et al. Regulated fluctuations in nanog expression mediate cell fate decisions in embryonic stem cells. PLoS Biol. 7, e1000149 (2009).

23. Filipczyk, A. et al. Network plasticity of pluripotency transcription factors in embryonic stem cells. Nat. Cell Biol. 17, 1235–1246 (2015).

24. Garg, A., Di Cara, A., Xenarios, I., Mendoza, L. & De Micheli, G.Synchronous versus asynchronous modeling of gene regulatory networks. Bioinforma. Oxf. Engl. 24, 1917–1925 (2008).

25. Albert, I., Thakar, J., Li, S., Zhang, R. & Albert, R.Boolean network simulations for life scientists. Source Code Biol. Med. 3, 16 (2008).

26. De Los Angeles, A. et al. Hallmarks of pluripotency. Nature 525, 469–478 (2015).

27. Kim, J., Chu, J., Shen, X., Wang, J. & Orkin, S. H.An Extended Transcriptional Network for Pluripotency of Embryonic Stem Cells. Cell 132, 1049–1061 (2008).

28. Chickarmane, V. & Peterson, C. A computational model for understanding stem cell, trophectoderm and endoderm lineage determination. PloS One 3, e3478 (2008).

29. Tam, W.-L. et al. T-cell factor 3 regulates embryonic stem cell pluripotency and selfrenewal by the transcriptional control of multiple lineage pathways. Stem Cells Dayt. Ohio 26, 2019–2031 (2008).

30. Bao, S. et al. Epigenetic reversion of post-implantation epiblast to pluripotent embryonic stem cells. Nature 461, 1292–1295 (2009).

31. Borgel, J. et al. Targets and dynamics of promoter DNA methylation during early mouse development. Nat. Genet. 42, 1093–1100 (2010).

32. Gillich, A. et al. Epiblast stem cell-based system reveals reprogramming synergy of germline factors. Cell Stem Cell 10, 425–439 (2012).

33. De Cegli, R. et al. Reverse engineering a mouse embryonic stem cell-specific transcriptional network reveals a new modulator of neuronal differentiation. Nucleic Acids Res. 41, 711–726 (2013).

34. Kushwaha, R. et al. Interrogation of a context-specific transcription factor network identifies novel regulators of pluripotency. Stem Cells Dayt. Ohio 33, 367–377 (2015).

35. Ma, S., Gong, Q. & Bohnert, H. J. An Arabidopsis gene network based on the graphical Gaussian model. Genome Res. 17, 1614-1625 (2007).

36. Kolodziejczyk, A. A. et al. Single Cell RNA-Sequencing of Pluripotent States Unlocks Modular Transcriptional Variation. Cell Stem Cell 17, 471–485 (2015).

37. Hayashi, K., Lopes, S. M. C. de S., Tang, F. & Surani, M.A. Dynamic equilibrium and heterogeneity of mouse pluripotent stem cells with distinct functional and epigenetic states. Cell Stem Cell 3, 391–401 (2008).

38. Brons, I. G. M. et al. Derivation of pluripotent epiblast stem cells from mammalian embryos. Nature 448, 191–195 (2007).

39. Ying, Q.-L. et al. The ground state of embryonic stem cell self-renewal. Nature 453, 519–523 (2008).

40. Marks, H. et al. The transcriptional and epigenomic foundations of ground state pluripotency. Cell 149, 590–604 (2012).

41. Bernemann, C. et al. Distinct developmental ground states of epiblast stem cell lines determine different pluripotency features. Stem Cells Dayt. Ohio 29, 1496–1503 (2011).

42. Festuccia, N. et al. Esrrb is a direct Nanog target gene that can substitute for Nanog function in pluripotent cells. Cell Stem Cell 11, 477–490 (2012).

43. Hanna, J. et al. Direct cell reprogramming is a stochastic process amenable to acceleration. Nature 462, 595–601 (2009).

44. Martello, G. et al. Esrrb is a pivotal target of the Gsk3/Tcf3 axis regulating embryonic stem cell self-renewal. Cell Stem Cell 11, 491–504 (2012).

45. Tai, C.-I. & Ying, Q.-L. Gbx2, a LIF/Stat3 target, promotes reprogramming to and retention of the pluripotent ground state. J Cell Sci 126, 1093–1098 (2013).

46. Joo, J. Y. et al. Establishment of a primed pluripotent epiblast stem cell in FGF4-based conditions. Sci. Rep. 4, 7477 (2014).

47. Wray, J. et al. Inhibition of glycogen synthase kinase-3 alleviates Tcf3 repression of the pluripotency network and increases embryonic stem cell resistance to differentiation. Nat. Cell Biol. 13, 838–845 (2011).

48. Yeo, J.-C. et al. Klf2 is an essential factor that sustains ground state pluripotency. Cell Stem Cell 14, 864–872 (2014).

49. Wray, J., Kalkan, T. & Smith, A. G.The ground state of pluripotency. Biochem. Soc. Trans. 38, 1027–1032 (2010).

50. Martello, G., Bertone, P. & Smith, A.Identification of the missing pluripotency mediator downstream of leukaemia inhibitory factor. EMBO J. 32, 2561–2574 (2013).

51. Smith, A. G. et al. Inhibition of pluripotential embryonic stem cell differentiation by purified polypeptides. Nature 336, 688–690 (1988).

52. ten Berge, D. et al. Embryonic stem cells require Wnt proteins to prevent differentiation to epiblast stem cells. Nat. Cell Biol. 13, 1070–1075 (2011).

53. Hayashi, Y. et al. BMP4 induction of trophoblast from mouse embryonic stem cells in defined culture conditions on laminin. In Vitro Cell. Dev. Biol. Anim. 46, 416–430 (2010).

54. Niwa, H. et al. Interaction between Oct3/4 and Cdx2 determines trophectoderm differentiation. Cell 123, 917–929 (2005).

55. Morgani, S. M. et al. Totipotent embryonic stem cells arise in ground-state culture conditions. Cell Rep. 3, 1945–1957 (2013).

56. Vallier, L. et al. Early cell fate decisions of human embryonic stem cells and mouse epiblast stem cells are controlled by the same signalling pathways. PloS One 4, e6082 (2009).

57. Bernardo, A. S. et al. BRACHYURY and CDX2 mediate BMP-induced differentiation of human and mouse pluripotent stem cells into embryonic and extraembryonic lineages. Cell Stem Cell 9, 144–155 (2011).

58. Rugg-Gunn, P. J. et al. Cell-surface proteomics identifies lineage-specific markers of embryo-derived stem cells. Dev. Cell 22, 887–901 (2012).

59. Morris, S. A. et al. Dissecting engineered cell types and enhancing cell fate conversion via CellNet. Cell 158, 889–902 (2014).

60. Herberg, M., Kalkan, T., Glauche, I., Smith, A. & Roeder, I.A model-based analysis of culture-dependent phenotypes of mESCs. PloS One 9, e92496 (2014).

61. Glauche, I., Herberg, M. & Roeder, I. Nanog variability and pluripotency regulation of embryonic stem cells--insights from a mathematical model analysis. PloS One 5, e11238 (2010).

62. Marucci, L. et al. β-catenin fluctuates in mouse ESCs and is essential for Nanog-mediated reprogramming of somatic cells to pluripotency. Cell Rep. 8, 1686–1696 (2014).

63. O’Malley, J. et al. High-resolution analysis with novel cell-surface markers identifies routes to iPS cells. Nature 499, 88–91 (2013).

64. Karwacki-Neisius, V. et al. Reduced Oct4 expression directs a robust pluripotent state with distinct signaling activity and increased enhancer occupancy by Oct4 and Nanog. Cell Stem Cell 12, 531–545 (2013).

65. Iovino, N. & Cavalli, G. Rolling ES cells down the Waddington landscape with Oct4 and Sox2. Cell 145, 815–817 (2011).

66. Yamanaka, S.Elite and stochastic models for induced pluripotent stem cell generation. Nature 460, 49–52 (2009).

67. Rayon, T. et al. Notch and hippo converge on Cdx2 to specify the trophectoderm lineage in the mouse blastocyst. Dev. Cell 30, 410–422 (2014).

68. Ng, R. K. et al. Epigenetic restriction of embryonic cell lineage fate by methylation of Elf5. Nat. Cell Biol. 10, 1280–1290 (2008).

69. Cambuli, F. et al. Epigenetic memory of the first cell fate decision prevents complete ES cell reprogramming into trophoblast. Nat. Commun. 5, 5538 (2014).

70. Qiao, W. et al. Intercellular network structure and regulatory motifs in the human hematopoietic system. Mol. Syst. Biol. 10, 741 (2014).

71. Meacham, C. E. & Morrison, S. J.Tumour heterogeneity and cancer cell plasticity. Nature 501, 328–337 (2013).

72. Heng, H. H. Q. et al. Genetic and epigenetic heterogeneity in cancer: a genome-centric perspective. J. Cell. Physiol. 220, 538–547 (2009).

73. Moignard, V. et al. Decoding the regulatory network of early blood development from single-cell gene expression measurements. Nat. Biotechnol. 33, 269–276 (2015).

74. Chang, K. H. & Zandstra, P. W.Quantitative screening of embryonic stem cell differentiation: endoderm formation as a model. Biotechnol. Bioeng. 88, 287–298 (2004).

75. Tanaka, S., Kunath, T., Hadjantonakis, A. K., Nagy, A. & Rossant, J.Promotion of trophoblast stem cell proliferation by FGF4. Science 282, 2072–2075 (1998).

76. Spandidos, A., Wang, X., Wang, H. & Seed, B. PrimerBank: a resource of human and mouse PCR primer pairs for gene expression detection and quantification. Nucleic Acids Res. 38, D792–799 (2010).

77. Hadjantonakis, A.-K. & Papaioannou, V. E. Dynamic in vivo imaging and cell tracking using a histone fluorescent protein fusion in mice. BMC Biotechnol. 4, 33 (2004).

